# Insulin-like growth factor 2 (IGF2) protects against Huntington’s disease through the extracellular disposal of protein aggregates

**DOI:** 10.1101/2020.05.28.119164

**Authors:** Paula García-Huerta, Paulina Troncoso-Escudero, Di Wu, Arun Thiruvalluvan, Marisol Cisternas, Daniel R. Henríquez, Lars Plate, Pedro Chana-Cuevas, Cristian Saquel, Peter Thielen, Kenneth A. Longo, Brad J. Geddes, Gerardo L. Lederkremer, Neeraj Sharma, Marina Shenkman, Swati Naphade, Lisa M. Ellerby, Pablo Sardi, Carlos Spichiger, Hans G. Richter, Felipe A. Court, R. Luke Wiseman, Christian Gonzalez-Billault, Steven Bergink, Rene L. Vidal, Claudio Hetz

## Abstract

Impaired neuronal proteostasis is a salient feature of many neurodegenerative diseases, highlighting alterations in the function of the endoplasmic reticulum (ER). We previously reported that targeting the transcription factor XBP1, a key mediator of the ER stress response, delays disease progression and reduces protein aggregation in various models of neurodegeneration. To identify disease-modifier genes that may explain the neuroprotective effects of XBP1 deficiency, we performed gene expression profiling of brain cortex and striatum of these animals and uncovered insulin-like growth factor 2 (*Igf2*) as the major upregulated gene. Here we studied the impact of IGF2 signaling on protein aggregation in models of Huntington’s disease (HD) as proof-of-concept. Cell culture studies revealed that IGF2 treatment decreases the load of intracellular aggregates of mutant huntingtin and a polyglutamine peptide. These results were validated using induced pluripotent stem cells (iPSC)-derived medium spiny neurons from HD patients. The reduction in the levels of mutant huntingtin was associated with a decrease in the half-life of the intracellular protein. The decrease in the levels of abnormal protein aggregation triggered by IGF2 were independent of the activity of autophagy and the proteasome pathways, the two main routes for mutant huntingtin clearance. Conversely, IGF2 signaling enhanced the secretion of soluble mutant huntingtin species through exosomes and microvesicles involving changes in actin dynamics. Administration of IGF2 into the brain of HD mice using gene therapy led to a significant decrease in the levels of mutant huntingtin in three different animal models. Moreover, analysis of human post-mortem brain tissue, and blood samples from HD patients showed a reduction of IGF2 level. This study identifies IGF2 as a relevant factor deregulated in HD, operating as a disease modifier that buffers the accumulation of abnormal protein aggregates.

**One sentence summary:** IGF2 reduces the load of intracellular protein aggregates through the extracellular disposal of the mutant protein.

## Introduction

Protein misfolding and aggregation are considered one of the initial events resulting in neurodegeneration in a variety of brain diseases including amyotrophic lateral sclerosis (ALS), Parkinson’s, Alzheimer’s, fronto-temporal dementia and Huntington’s disease (HD), which are collectively classified as protein misfolding disorders (PMDs) (*1, 2*). The proteostasis network is fundamental to maintain the stability of the proteome and prevent abnormal protein aggregation (*3*). Chaperones resident in the cytosol and the endoplasmic reticulum (ER) ensure precise folding of newly synthesized native proteins. Quality control mechanisms identify misfolded proteins and facilitate their removal via the proteasome, lysosome and autophagy pathways (*4*). Importantly, the buffering capacity of the proteostasis network is reduced during aging, which may increase the risk to accumulate protein aggregates and undergo neurodegeneration (*4, 5*). One of the main nodes of the proteostasis network altered in neurodegenerative diseases involves the ER, the main site for protein folding and quality control in the cell (*6–8*). ER stress instigates a rapid and coordinated signaling pathway known as the unfolded protein response (UPR) to enforce adaptive programs and restore proteostasis (*9*). Conversantly, irremediable ER stress results in cell death by apoptosis (*10*).

The UPR is initiated by the activation of specialized stress transducers highlighting inositol-requiring transmembrane kinase/endonuclease (IRE1α) as the most conserved signaling branch (*11*). IRE1α is an ER-located kinase and endoribonuclease that, upon activation, controls the processing of the mRNA encoding X-box-binding protein 1 (XBP1), resulting in the expression of a stable and active transcription factor known as XBP1s (*12*). XBP1s upregulates genes related to protein folding, quality control, ER translocation, ERAD, among other processes (*13–15*). In *C. elegans* organisms, the expression of XBP1 in neurons controls organism aging (*16*), which may involve a crosstalk with classical signaling pathways regulating normal aging including the insulin-like growth factor (IGF)-1 and FOXO (DAF16) pathways (*17*). We have previously studied the functional contribution of XBP1 to various neurodegenerative conditions. Despite initial expectations that targeting this central UPR mediator will exacerbate disease progression, we observed that genetic disruption of the IRE1α/XBP1 pathway in the brain led to neuroprotection in models of ALS (*18*), Parkinsońs disease (*19*), Alzheimeŕs disease (*20, 21*) and HD (*22*), associated with reduced accumulation of abnormal protein aggregates. These findings illustrate the broader significance of XBP1 and the UPR to the progression of neurodegenerative diseases, highlighting the need to identify the mechanisms of action mediating neuroprotection.

To uncover novel regulatory elements that may underlie the beneficial effects of ablating XBP1 expression in the brain, here we performed a gene expression profile study in cortex and striatum of a XBP1 conditional knockout mice (XBP1cKO) and identified *Igf2* as the major upregulated gene. We then investigated the significance of IGF2 to abnormal protein aggregation using HD models as proof-of concept. HD is an autosomal disease, which belongs to a group of tandem repeat diseases caused by a CAG codon expansion known as polyglutamine disorders. HD is caused by mutations in the huntingtin (Htt) gene, involving the generation of an abnormal tract of over 40 glutamine residues (polyQ) at the N-terminal region (*23*). HD patients develop a combination of motor, cognitive and psychological symptoms, associated with progressive neurodegeneration of the striatum and cerebral cortex (*24*). Mutant Htt (mHtt) often forms insoluble inclusions that alter cellular homeostasis including the function of the ER (*25, 26*). Here, we provide evidence indicating that IGF2 signaling decreases the accumulation of intracellular mHtt and polyglutamine peptide aggregates in various cell culture models of HD. Unexpectedly, the decrease of mutant huntingtin and polyglutamine peptide aggregates was independent of the activity of the proteasome or autophagy pathways. Instead, IGF2 treatment enhanced the non-conventional disposal of soluble polyglutamine peptide species into the extracellular space through the secretion via microvesicles and exosomes. To validate our results *in vivo*, we developed a therapeutic approach to deliver IGF2 into the brain using recombinant viruses, which resulted in significant reduction of mHtt levels in three different preclinical models of HD. Furthermore, we observed a dramatic reduction in IGF2 levels in brain and blood samples derived from HD patients, in addition to medium spiny neurons generated from HD iPSCs. We propose that IGF2 signaling operates as a disease modifier, where its deregulation enhances the abnormal accumulation of mHTT in HD. Overall, this study uncovered a previously unanticipated role of IGF2 in handling protein aggregates in HD.

## Results

### Igf2 is upregulated in the brain of XBP1 deficient mice

To define the regulatory network involved in the neuroprotective effects triggered by XBP1 deficiency in the brain, we generated XBP1_cKO_ mice using the Nestin-Cre system (*27*). Animals of 12 to 14 months old were euthanized and brain tissue was dissected to obtain both cortex and striatum to perform a global gene expression profile analysis using a total of 29 animals. Surprisingly, global analysis indicated that the extent of gene expression changes was modest. A comparison between the two brain tissues indicated poor overlap in gene expression changes between brain cortex and striatum where only the expression of *Igf2* was modified in both brain areas, showing higher expression in XBP1_cKO_ mice (Fig. 1A).

**Figure 1.**
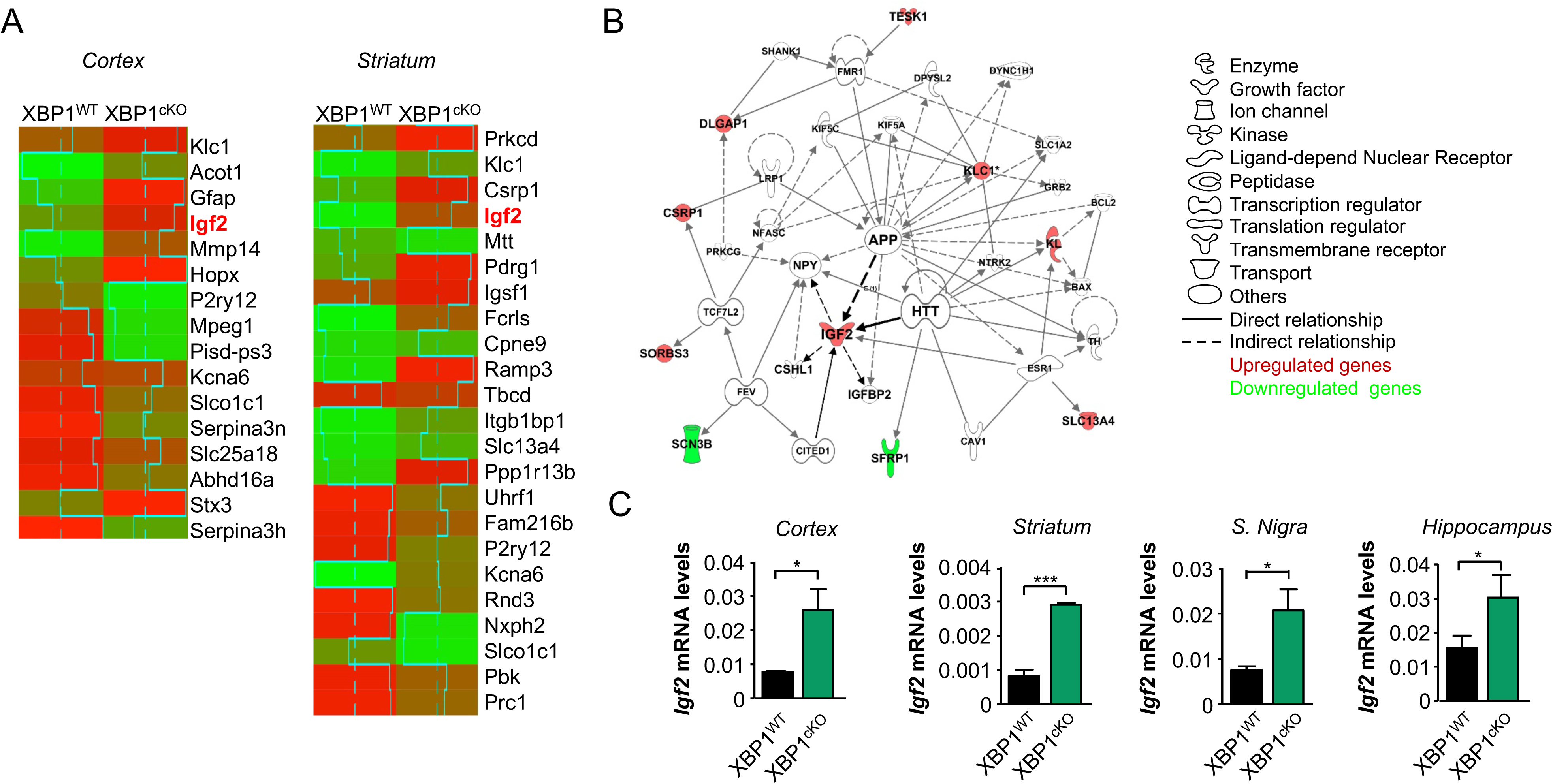
XBP1 deficiency increases *Igf2* expression levels. **(A)** Gene expression was determined in the brain cortex and striatum of XBP1_cKO_ (n = 17) and litter mate control (n = 12) animals using Illumina BeadChip® platform and analyzed as described in materials and methods. Heatmaps of microarray analysis of cortex or striatum tissue of two experimental groups considering 1.5 or 1.6-fold change as cutoff, respectively. The normalized average signal of each group was displayed in the heatmap. Red color is used to indicate upregulated genes and green color for downregulation. **(B)** Network of significantly enriched in genes differentially expressed in the brain cortex and striatum of XBP1_cKO_. This set of genes was grouped in the Ingenuity Top Network 2 ‘Cancer, Drug Metabolism, Endocrine System Development and Function’. Red: significantly up-regulated gene; Green: significantly down-regulated gene; Gray: non-significant differential expression; White: biological terms related with the network. **c** *Igf2* mRNA levels were measured by real-time PCR in cDNA generated from cortex, striatum, substantia nigra and hippocampus region of 9-months old XBP_WT_ and XBP1_cKO_ mice. Data represents the average and SEM of the analysis of 3-6 animals per group. Statistically significant differences detected by one-tailed unpaired *t*-test (***: *p* < 0.001; *: *p* < 0.05).

Although the top regulated genes in XBP1_cKO_ mice displayed a consistent fold change of at least 1.5-fold, few genes including *Igf2* showed a robust statistical significantly (q-value = 0) (Supplementary Table S1). Ingenuity pathway analysis (IPA) suggested a relation between *Igf2* with key biological pathways related to neuronal dysfunction associated with Htt and the amyloid precursor protein (Fig. 1B). A second IPA analysis found Htt as candidate of “upstream regulators” with several target genes deregulated (*Igf2, Gfap, Mmp14, Mpeg1* and *Serpine3*) and lowest p-value (data no shown).

We then validated the changes in the expression levels of *Igf2* in the brain of XBP1_cKO_ mice using real-time PCR and detected a significant increase when compared to wild-type littermates (Fig. 1C). To determine whether the upregulation of *Igf2* occurs in the context of neurodegenerative diseases when XBP1 expression is ablated, we analyzed brain tissue derived from a HD model (YAC128 transgenic mice) that was crossed with our XBP1_cKO_ mice (*22*). We measured mRNA levels in the striatum of XBP1_cKO_-YAC128 mice and confirmed the increase in the levels of *Igf2* compared to control animals (Supplementary Fig. S1A). As control, we also monitored *Igf1* levels and did not detect any significant differences between XBP1_cKO_-YAC128 and control mice (Supplementary Fig. S1B), indicating that *Xbp1* deficiency in the nervous system results in the selective upregulation of *Igf2* expression.

### IGF2 reduces polyQ and mHtt aggregation

Although IGF2 is the less studied member of the insulin-like peptide family, which includes IGF1 and insulin, recent studies have uncovered the importance of IGF2 in brain physiology and neurodegeneration. For example, IGF2 overexpression restored memory function and decreased the accumulation of amyloid β in models of AD (*28, 29*). In ALS, IGF2 is differentially expressed in resistant motoneurons and its overexpression extends the survival of mutant SOD1 mice (*30*). However, the mechanisms involved in these protective activities needs to be further defined. Since the accumulation of misfolded protein in neurons is a common feature of all of these diseases, we decided to explore the possible function of IGF2 as a disease modifier agent in PMDs.

We first performed cell culture studies to determine the consequences of IGF2 expression over abnormal protein aggregation. Neuro2a cells were transfected with expression vectors for polyQ_79_-EGFP together with IGF2 or empty vector (Mock) followed by the analysis of intracellular inclusions. Quantification of the number of cells containing GFP puncta by fluorescence microscopy revealed a strong reduction in the number of polyQ_79_-EGFP inclusions, when co-transfected with IGF2 vector (Fig. 2A). Then, we monitored the aggregation of polyQ_79_-EGFP using biochemical approaches. Again, IGF2 or IGF2-HA expression led to a near complete reduction in the accumulation of both high molecular weight and detergent insoluble and monomeric species of polyQ_79_-EGFP using standard detection by western blot (Fig. 2B and Supplementary Fig. 2A, respectively), whereas polyQ_79_-EGFP mRNA levels were equal in both conditions (Supplementary Fig. 2B). Similar results were obtained when protein extracts were analyzed by filter trap, an assay that detects protein aggregates by size using a 200 nm cellulose acetate filter (Fig. 2C and Supplementary Fig. S2C). Because similar results were obtained with both IGF2 and IGF2-HA constructs, the following experiments were performed using the tagged version.

**Figure 2.**
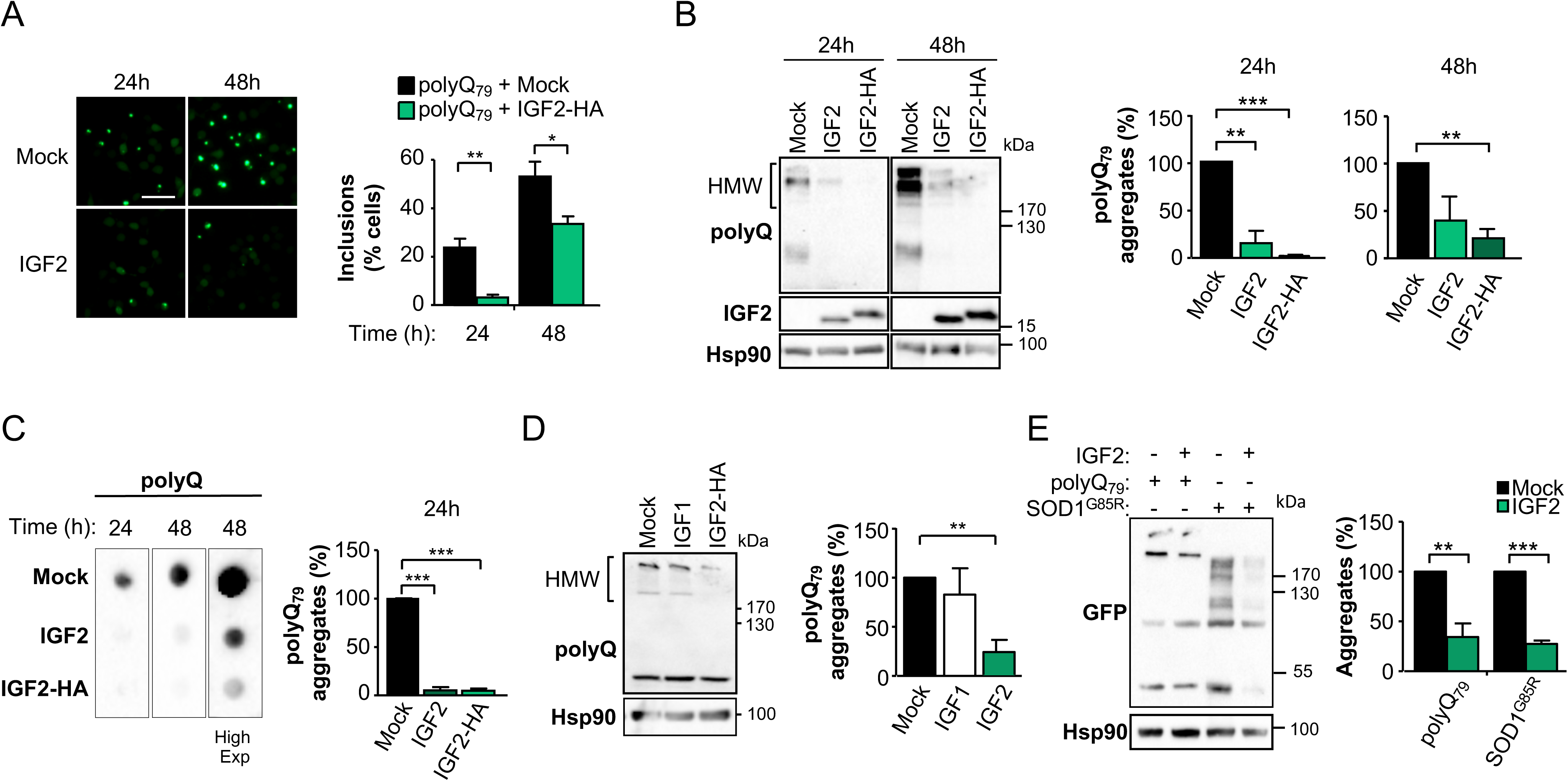
IGF2 expression reduces the levels of polyQ_79_ aggregates. **(A)** Neuro2a cells were co-transfected with a polyQ_79_-EGFP expression vector and IGF2 plasmid or empty vector (Mock). 24 or 48 h later, the number of polyQ_79_-EGFP inclusions were visualized by fluorescence microscopy (left panel). Quantification of inclusions was normalized by the number of GFP-positive cells (right panel; n = 100 to 350 cells per experiment). Scale bar, 20 µm. **(B)** PolyQ_79_-EGFP expression was analyzed in whole cell extracts by western blot analysis using an anti-GFP antibody. A non-tagged version of IGF2 expression vector was also included. The presence of high molecular weight (HMW) species is indicated. Hsp90 and IGF2 levels were also determined (left panel). PolyQ_79_-EGFP aggregates levels were quantified and normalized to Hsp90 levels (right panel). **(C)** Filter trap assay was performed in the same cells extracts analyzed in (B) (left panel) and quantified at 24 h (right panel). **(D)** Neuro2a cells were transiently co-transfected with polyQ_79_-EGFP together with plasmids to express IGF1, IGF2 or empty vector (Mock). The presence of polyQ_79_-EGFP aggregates was analyzed in whole cell extracts after 24 h by western blot analysis. Hsp90 expression was monitored as loading control. **(E)** Neuro2a cells were co-transfected with polyQ_79_-EGFP or SOD1_G85R_-GFP and IGF2 plasmid or empty vector (Mock). After 24 h, polyQ_79_ and SOD1 aggregation were measured in total cell extracts using western blot under non-reducing conditions (without DTT). Hsp90 expression was monitored as loading control. In all quantifications, values represent the mean and SEM of at least three independent experiments. Statistically significant differences detected by two-tailed unpaired *t*-test (***: *p* < 0.001; **: *p* < 0.01; *: *p* < 0.05).

To evaluate if the effects on aggregation were specific for IGF2, we measured aggregation levels of polyQ_79_-EGFP in cells co-transfected with an IGF1 expression vector. Side-by-side comparison indicated that only IGF2 reduced the levels of polyQ_79_-EGFP aggregates (Fig. 2D). Since XBP1 deficiency also affects the accumulation of protein aggregates associated with other diseases, we performed additional experiments to determine the effects of IGF2 over ALS-linked mutant SOD1_G85R_. Remarkably, the co-expression of mutant SOD1 with IGF2-HA also modified its aggregation as assessed by western blot, in addition to reduce the levels of its monomeric form, similar to the effects observed with polyQ_79-_EGFP aggregation (Fig. 2E), suggesting broader implications to neurodegeneration.

To define if IGF2 exerts its function from the extracellular compartment, we treated cells expressing polyQ_79_-EGFP with conditioned media enriched in IGF2. In these experiments we also included a construct that expresses a fragment of mutant huntingtin spanning exon 1 with 85 CAG repeats (GFP-mHttQ_85_). We transiently expressed polyQ_79_-EGFP or GFP-mHttQ_85_ in Neuro2a cells pre-treated with IGF2-enriched media generated from Neuro2a cells transfected with IGF2 or empty vector for 24 h. A clear reduction in the extent of protein aggregation was observed in cells exposed to IGF2-enriched media using three different assays (Fig. 3A-C). These results suggest that IGF2 may exert its function in a paracrine manner, probably activating signaling pathways through the binding to a membrane receptor. Interestingly, treatment of cells with insulin did not reduce polyQ_79_-EGFP aggregation levels in our cellular model (Fig. 3D), similar to the results obtained with IGF1 overexpression.

**Figure 3.**
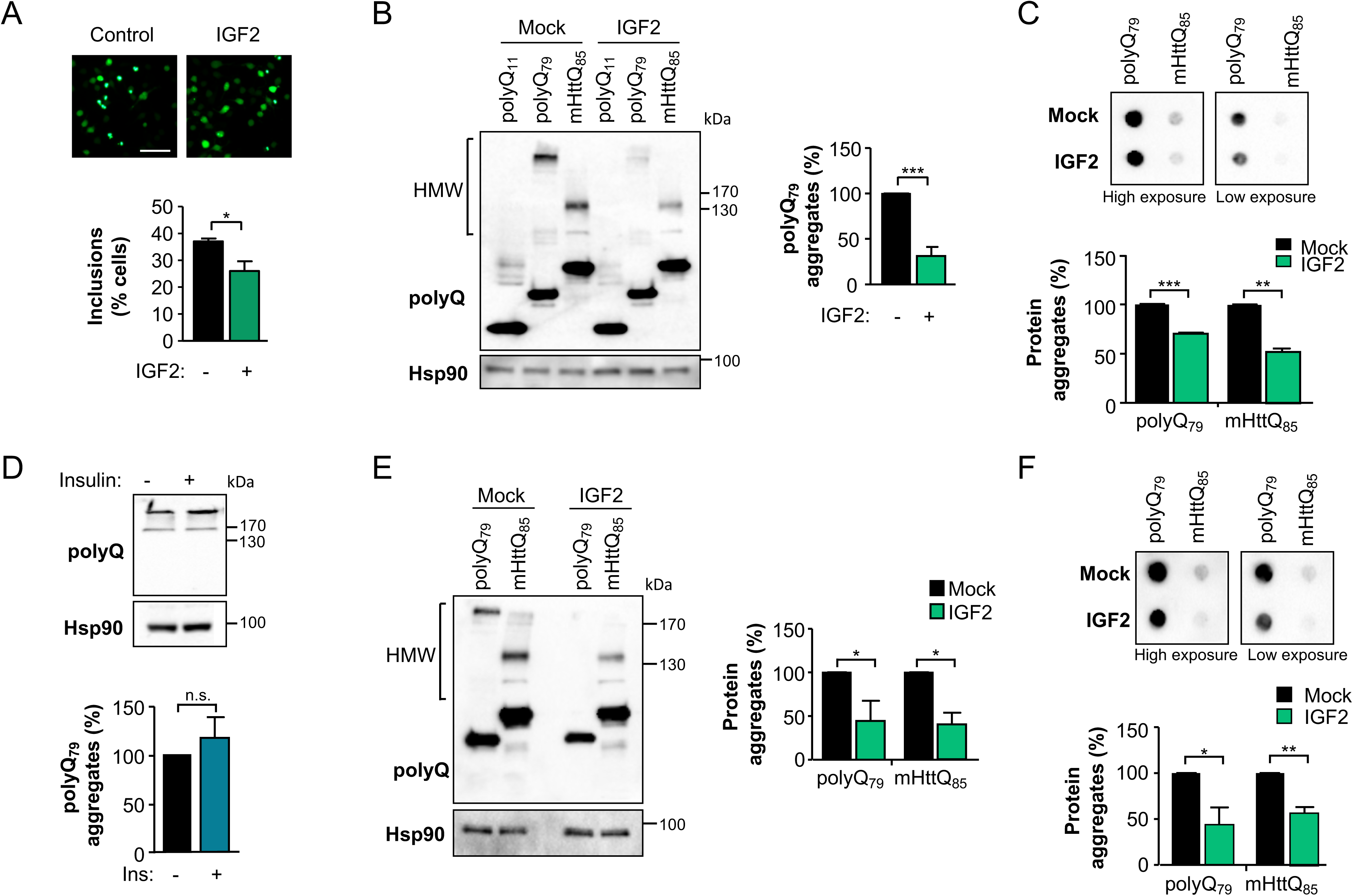
IGF2 treatment reduces the accumulation of polyQ_79_ and mHttQ_85_ aggregates. **(A)** Neuro2a cells were transfected with polyQ_79_-EGFP expression vectors in the presence of IGF2-enriched cell culture media. 24 h later polyQ_79_-EGFP and mHttQ_85_-GFP inclusions were visualized by microscopy (upper panel) and quantified (bottom panel; n = 100 to 350 cells per experiment). Scale bar, 20 µm. **(B)** Neuro2a cells were transfected with polyQ_79_-EGFP or mHttQ_85_-GFP expression vectors in presence of IGF2-enriched cell culture media. 24 h later the aggregation levels of polyQ_79_-EGFP or mHttQ_85_-GFP were determined by western blot analysis using anti-GFP antibody. Hsp90 expression was monitored as loading control. **(C)** Filter trap assay was performed using the same protein extracts analyzed in (b) to quantify polyQ_79_-EGFP and mHttQ_85_-GFP aggregates. **(D)** Neuro2a cells were transfected with polyQ_79_-EGFP in presence of 1 μM insulin. 24 h later the aggregation of polyQ_79_-EGFP protein was measured in total cell extracts by western blot. Hsp90 expression was monitored as loading control. **(E)** Neuro2a cells were transfected with polyQ_79_-EGFP or mHttQ_85_-GFP expression vectors. After 24 h cells were treated with IGF2-enriched cell culture media and 24 h later aggregation levels determined and quantified by western blot analysis using anti-GFP antibody. Hsp90 levels were monitored as loading control. **(F)** Filter trap assay was performed in the same cell extracts analyzed in (e) and polyQ_79_-EGFP and mHttQ_85_-GFP aggregates were detected using anti-GFP antibody. In all quantifications, average and SEM of at least three independent experiments are shown. Statistically significant differences detected by two-tailed unpaired *t*-test (***: *p* < 0.001; **: *p* < 0.01; *: *p* < 0.05).

We then evaluated if IGF2 could reduce the content of pre-formed aggregates. Thus, we expressed polyQ_79_-EGFP or GFP-mHttQ_85_ for 24 h and then treated Neuro2a cells with IGF2-enriched or control media. In agreement with our previous results, exposure of cells to IGF2-conditioned media attenuated the load of misfolded polyQ_79_-EGFP or GFP-mHttQ_85_ species (Fig. 3E, F), suggesting that IGF2 may trigger the degradation or disaggregation of abnormal protein aggregates.

### IGF2 reduces the aggregation of expanded polyglutamine proteins

To further validate our results, we evaluated the effects of IGF2 expression on a cellular model that expresses full-length mHtt. To this aim, HEK293 cells were transfected with expression vectors for a myc-tagged version of full-length mHtt with a repetition of 103 glutamines (FL HttQ_103-_myc) together with IGF2 or empty vector (Mock). Again, we observed that the expression of IGF2 caused a significant reduction in the levels of both monomeric and detergent insoluble species of the FL HttQ_103_-myc using western blot analysis (Fig. 4A). These results were also confirmed when cell lysates were analyzed by dot blot (Fig. 4B). Overall, IGF2 expression leads to decrease levels of monomeric and aggregated forms of full-length mHtt.

**Figure 4.**
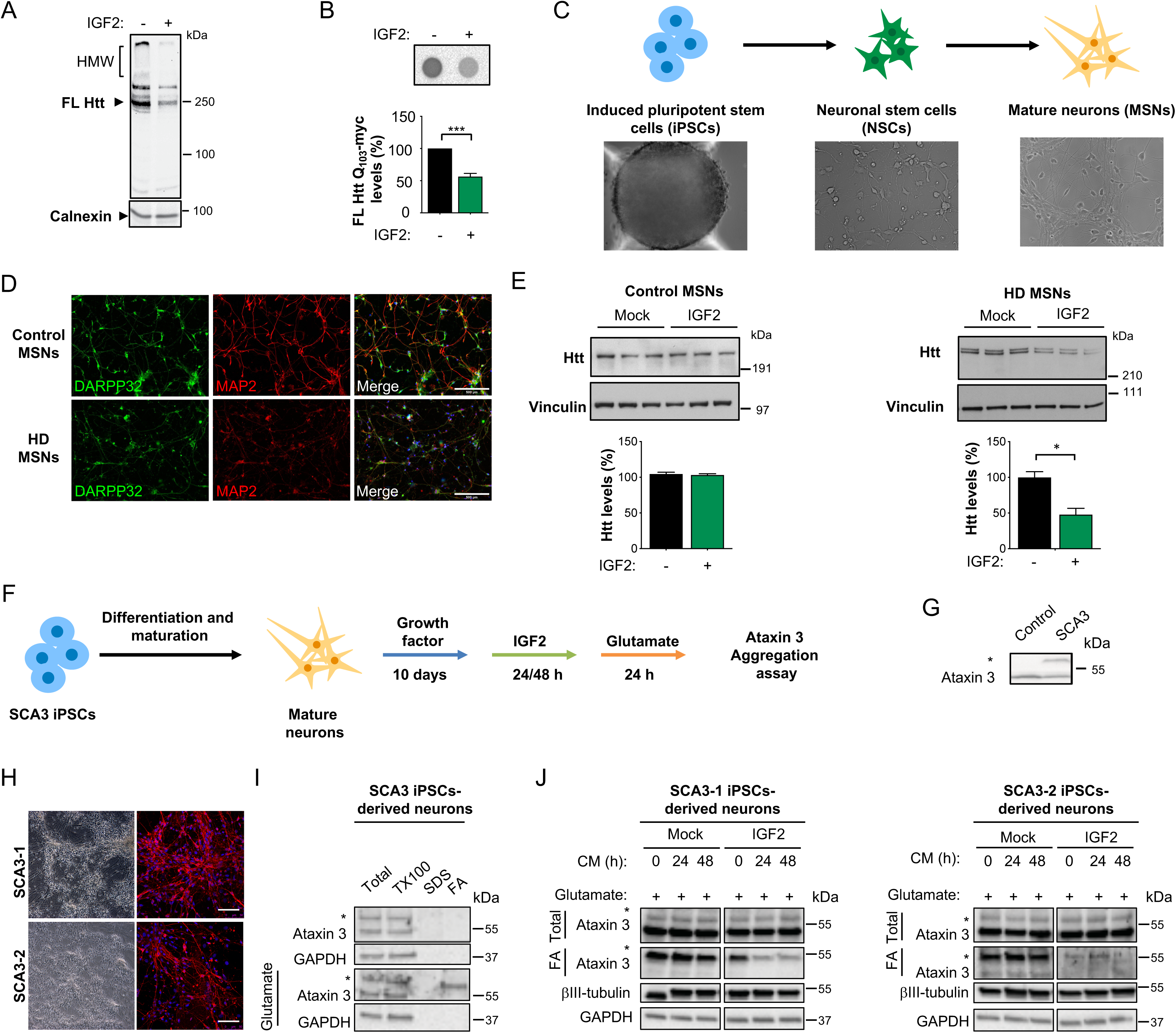
IGF2 overexpression reduced the levels of huntingtin in MSNs derived from HD patients. **(A)** HEK293T cells were co-transfected with full length (FL) HttQ_103_-myc and IGF2 or empty vector (Mock). 24 h later, FL HttQ_103_-myc levels were analyzed in whole cell extracts by western blot analysis using an anti-Htt antibody. The arrow indicates the migration of full-length huntingtin. **(B)** HEK293T cells were co-transfected with FL HttQ_103_-myc and IGF2 or empty vector (Mock). After 24 h, FL HttQ_103_-myc was detected in cell lysates by dot blot. Statistically significant differences detected by two-tailed unpaired *t*-test (***: *p* < 0.001). **(C)** Schematic representation of the differentiation process to generate mature medium spiny neurons (MSNs) from induced human pluripotent stem cells (iPSCs), showing cell morphology using contrast phase microscopy. **(D)** Confirmation of the expression of GABAergic (DARPP-32) and neuronal (MAP2) markers in control and HD patients MSNs using immunofluorescence. **(E), (F)** MSNs from control subjects and HD patients were transduced with adeno-associated viral (AAV) vectors expressing IGF2 o empty vector (Mock). 17 days later, huntingtin protein levels were measured in total protein extracts by western blot using anti-Htt antibody. Vinculin expression was monitored as loading control (lower panel). Values represent the mean and SEM of three independent experiments. Statistically significant differences detected by one-way ANOVA (*: *p* < 0.05). **(G)** Schematic representation of the generation of neuronal cultures generated from SCA-derived iPSCs. **(H)** Ataxin 3 levels were monitored in control and SCA3 patient fibroblasts using western blot analysis. * represent the mutant protein. **(I)** Differentiated SCA3 iPSC-derived neurons at 90 days post differentiation were stained for βIII-tubulin neuronal marker. Scale bars: 50 µm. **(J)** Neurons were treated with 0.1 mM L-glutamate for 1 h. Then, detergent insoluble protein extracts were generated in triton X100 (TX-100) and pellets were resuspended in SDS to yield an SDS-soluble fraction (SDS) and SDS-insoluble fraction that was then dissolved in formic acid (FA). Ataxin 3 levels were monitored in all fractions using western blots analysis (*-expanded allele). Two independent experiments were performed. **(K)** SCA3 cells were stimulated with IGF2-conditioned media for indicated time points followed by stimulation with 0.1 mM L-glutamate followed by western blot analysis as described in j. GAPDH levels were determined as loading control and βIII-tubulin as a neuronal marker.

Next, to increase the relevance of the current study, we moved forward and determined the possible effects of IGF2 in a human model of HD that do not rely on overexpression using induced pluripotent stem cells (iPSCs) derived from patients (*31, 32*). Thus, we generated human medium spiny neurons (MSNs) using iPSC-derived from HD patients and control subjects (Fig. 4C, D). To express IGF2, we packed its cDNA into an adeno-associated viral vector (AAV) using serotype 2, which has a high tropism for neurons. MSN cultures were transduced with AAV-IGF2 or AAV-Mock for 17 days followed by western blot analysis to determine the levels of mHtt. Remarkably, expression of IGF2 resulted in a decrease in the total levels of mHtt in HD-derived MSNs (Fig. 4E, right panel). Importantly, treatment of MSN cultures from control subjects did not modify the levels of wild-type Htt (Fig. 4E, left panel). Taken together, these results indicate that IGF2 expression reduces the content of mHtt in various models of HD.

The expansion of polyglutamine tracks is the underlying cause of different diseases, including six spinocerebellar ataxias (SCA), where SCA3 is the most common form of the disease, and the second form of polyglutamine disease after HD (*33*). SCA3 involves the expansion in the polyglutamine track in Ataxin-3, resulting in its abnormal protein aggregation. To validate the possible effects of IGF2 expression on the levels of other disease-related genes, we generated neuronal cultures using iPSC obtained from SCA3 patients (Fig. 4F-H) and exposed them to IGF2-enrriched conditioned cell culture media. In this experimental setting, exposure of cells to glutamate results in the abnormal aggregation of mutant Ataxin-3, reflected in the accumulation of the mutant protein in the detergent insoluble fraction (*34*) (Fig. 4I). Exposure of cells to IGF2-enriched media led to reduced levels of mutant Ataxin-3 aggregates (Fig. 4J). Thus, IGF2 has broader effects in reducing the abnormal aggregation of polyglutamine-containing proteins.

### IGF2 reduces the half-life of intracellular soluble mHtt

The reduction in the steady-state levels of mHtt aggregates induced by the exposure of cells to IGF2 may be explained by an attenuation in the synthesis rate or an increase its degradation rate. We performed pulse-label experiments in HEK293T cells co-transfected with a mHttQ_43_-GFP construct together with IGF2 or empty vector (Mock). To evaluate translation rate, a pulse with S_35_ labeled methionine and cysteine was performed for increasing intervals on a total period of 60 minutes. These experiments indicated that the biosynthesis of mHtt was identical in cells expressing or not IGF2 (Fig. 5A). In addition, general protein synthesis was not affected by IGF2 expression (Supplementary Fig. S3A). Then, we monitored the decay in the levels of mHttQ_43_-GFP in cell extracts using pulse-chase to define its half-life. Remarkably, expression of IGF2 dramatically reduced the half-life of mHttQ_43_-GFP in the cell extract (Fig. 5B), without affecting the global stability of the proteome (Supplementary Fig. S3B). In these experiments, we did not observe the presence of high-molecular weight species of mHttQ_43_, suggesting that IGF2 was affecting the stability of the soluble or monomeric forms. Taken together, our data suggests that IGF2 signaling reduces the content of total mHtt associated with a shortening of the half-life of the protein inside the cell.

**Figure 5.**
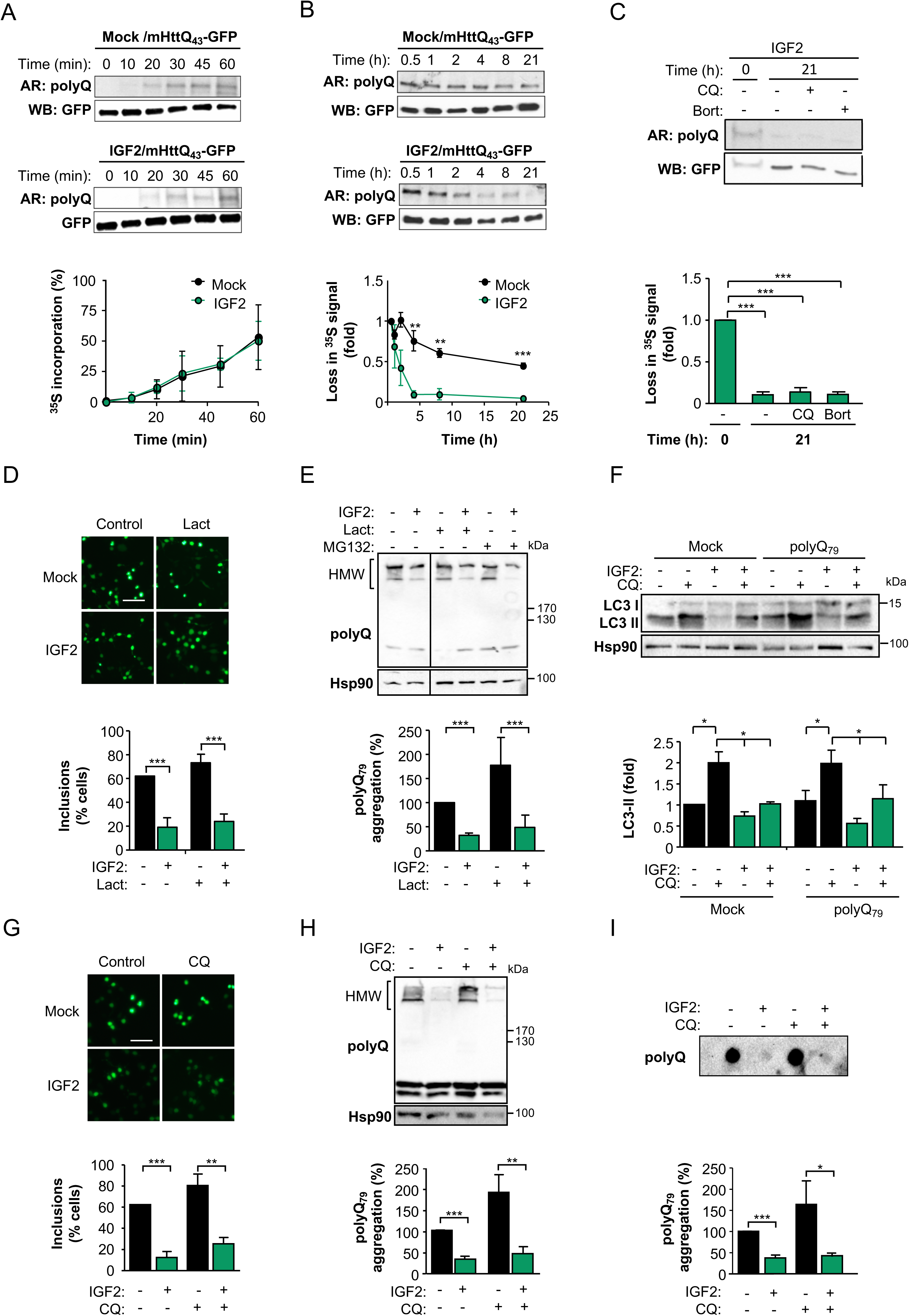
Stimulation of cells with IGF2 reduces the half-life of mHtt aggregates. **(A)** HEK293T cells were co-transfected with GFP-mHttQ_43_ expression vector with IGF2 plasmid (middle panel) or empty vector (Mock) (upper panel), and then pulse labeled with _35_S for indicated time points. Autoradiography (AR) indicated the _35_S signal for each time point. Data were quantified and normalized to time point 0 h (lower panel). **(B)** HEK293T cells were co-transfected with GFP-mHttQ_43_ expression vector with IGF2 plasmid (middle panel) or empty vector (Mock) (upper panel), and then pulse labeled with _35_S for indicated time points and the decay of the radioactive signal during chasing was monitored as described in materials and methods. Autoradiography (AR) indicated the _35_S signal for each time point. Data were quantified and normalized to the time point 1 h (lower panel). **(C)** HEK293T cells were transfected with GFP-mHttQ_43_ and IGF2 expression vectors. Pulse was performed 24 h after transfection. Cells were treated with 30 μM chloroquine (CQ) or 1 µM bortezomib (Bort) at the beginning of the chasing for additional 21 h (upper panel). Data were quantified and normalized to time point 1 h (lower panel). **(D)** Neuro2a cells were co-transfected with a polyQ_79_-EGFP expression vector and IGF2 plasmid or empty vector (Mock) for 8 h and then treated with 1 μM lactacystin (Lact) for additional 16 h (upper panel). Quantification of aggregates per GFP-positive cell was performed (lower panel; N=100 to 350 cells per experiment). **(E)** Neuro2a cells were co-transfected with a polyQ_79_-EGFP expression vector and IGF2 plasmid or empty vector (Mock) for 8 h, and then treated with 1 μM lactacystin (Lact) or 1 µM MG132 for additional 16 h. PolyQ_79_-EGFP aggregation was analyzed in whole cells extracts by western blot using anti-GFP antibody and quantified (lower panel). Hsp90 expression was analyzed as loading control (upper panel). **(F)** Neuro2a cells where co-transfected with a polyQ_79_-EGFP expression vector and IGF2 plasmid or empty vector (Mock) for 8 h, and then treated with 30 μM chloroquine (CQ) for additional 16 h. Endogenous lipidated LC3-II levels were monitored by western blot using anti-LC3 antibody. Hsp90 expression was monitored as loading control (upper panel). LC3 II levels were quantified and normalized to Hsp90 (lower panel). **(G)** Neuro2a cells were co-transfected with polyQ_79_-EGFP and IGF2 expression vectors or empty vector (Mock) for 8 h, and then treated with 30 μM CQ for additional 16 h (upper panel). Quantification of aggregates per GFP-positive cell was performed (lower panel; N=100 to 350 cells per experiment). **(H)** Neuro2a cells were co-transfected with polyQ_79_-EGFP and IGF2 expression vectors or empty vector (Mock) for 8 h, and then treated with 30 μM CQ for additional 16 h. PolyQ_79_-EGFP aggregation was analyzed in whole cells extracts by western blot using anti-GFP antibody and quantified (lower panel). Hsp90 expression was monitored as loading control (upper panel). **(I)** Filter trap was performed using the same cells extracts analyzed in (H). In all quantifications, average and SEM of at least three independent experiments are shown. Statistically significant differences detected by two-tailed unpaired *t*-test (***: *p* < 0.001; **: *p* < 0.01; *: *p* < 0.05).

### The autophagy and proteasome pathways are not involved in the reduction of polyQ aggregates induced by IGF2

Two major protein degradation routes are well established mediators of mHtt clearance, the proteasome and macroautophagy pathways (*35, 36*). We monitored mHtt levels after inhibiting lysosomal function with chloroquine or the proteasome with bortezomib. To this end, HEK293T cells were co-transfected with mHttQ_43_-GFP and IGF2 expression vectors followed by a radioactive pulse 24 h later to perform a chase experiment in the presence of the inhibitors. Surprisingly, neither chloroquine nor bortezomib treatments reverted the effects of IGF2 on mHtt levels (Fig. 5C). As controls for the activity of the inhibitors, we monitored the levels of LC3 and p62 (markers of autophagy-mediated degradation) and the pattern of total ubiquitination (Supplementary Fig. S3c).

Since the pulse chase methodology demonstrated that IGF2 decreases mHtt intracellular content, we performed further experiments to determine if autophagy or the proteasome mediate the reduction in the load of polyQ aggregates. Although a slight increase in the number of intracellular inclusions was observed at the basal condition after treating cells with the proteasome inhibitors lactacystin or MG132, no changes were observed in the number of polyQ_79_-EGFP inclusions in IGF2 expressing cells in presence of the inhibitors when compared to vehicle (Fig. 5D and Supplementary Fig. S4A). We confirmed these observations in Neuro2a cells by measuring polyQ_79_-EGFP aggregation by western blot and filter trap analysis (Fig. 5E and Supplementary Fig. S4B; see positive controls of total ubiquitination in Supplementary Fig. S4C). Again, the basal increase of polyQ aggregates after inhibiting the proteasome was lost in IGF2 treated cells. These results suggest that (i) IGF2 signaling may change the proportion between aggregate/soluble forms of polyQ inside the cell, or (ii) it may redirect polyQ aggregates toward a different pathway to mediate its clearance.

Based on our previous studies linking XBP1 deficiency with the upregulation of autophagy in neurons (*22*), we evaluated autophagy activation in cells overexpressing IGF2. Neuro2a cells were transfected with IGF2 in the presence or absence of polyQ_79_-EGFP expression vector. Then, autophagy was monitored after inhibiting lysosomal activity with chloroquine, followed by the analysis of LC3 and p62 levels (*37*). We observed that IGF2 did not stimulate autophagy in Neuro2a cells, but slightly reduced LC3-II accumulation (Fig. 5F). A similar trend was observed when we measured p62 levels in cells expressing polyQ_79_-EGFP (Supplementary Fig. S4D). Likewise, treatment of cells with chloroquine did not recover protein aggregation of cells exposed to IGF2 as measured with three independent methods (Fig. 5G-I). As control, in these experiments we monitored the accumulation of LC3-II and p62 in cells exposed to chloroquine (Supplementary Fig. S4E). Similarly, inhibition of lysosomal activity with chloroquine did not recover the normal levels of polyQ_79_-EGFP aggregation in HEK293T cells stimulated with IGF2 (Supplementary Fig. S4F, G, see positive controls in Supplementary Fig. S4H), confirming our previous results. Finally, we obtained similar results using the autophagy inhibitors bafilomycin A_1_ and 3-methyladenine (Supplementary Fig. S4I, J, respectively). Overall, these results suggest that the reduction in the load of polyQ_79_-EGFP aggregates induced by IGF2 treatment is independent of the proteasome and autophagy/lysosomal pathways. Our data suggests that IGF2 may reduce the levels of soluble and monomeric forms of mHtt or polyQ, reducing their aggregation levels by shifting the intracellular equilibrium toward a non-aggregated form. Alternatively, IGF2 signaling may reroute the clearance of protein aggregates toward an alternative pathway that is independent of the activity of the proteasome and macroautophagy/lysosomes.

### IGF2 signaling triggers polyQ secretion through extracellular vesicles

Since IGF2 administration bypassed the basal clearance of polyQ aggregates by the main degradation pathways, we explored the possibility that polyQ peptides are redirected to another compartment. mHtt has been found in the cerebrospinal fluid (*38*), as well as in neuronal allografts transplanted into the brain of HD patients (*39*) and can be secreted and transmitted from cell to cell. Recent studies in cell culture models validated this concept suggesting that mHtt can be exported to the extracellular space by non-conventional mechanisms (*40*). To determine the impact of IGF2 treatment in mHtt secretion, we monitored its presence in the cell culture media. Remarkably, enhanced secretion of polyQ_79_-EGFP was observed in cells expressing IGF2 when cell culture media was analyzed using dot blot (Fig. 6A, left panel). This phenomenon was also observed in cells overexpressing FL HttQ_103_-myc when IGF2 was expressed (Supplementary Fig. S4K). Similarly, analysis of SOD1_G85R_ levels in the cell culture media indicated that IGF2 induces its secretion (Fig. 6A, middle panel), whereas no changes where observed in a cytosolic marker such as tubulin (Fig. 6A, right panel), suggesting no cell lysis or cytosol leaking. In agreement with this, analysis of cell viability using propidium iodide staining and FACS analysis in cells expressing polyQ_79-_EGFP in the presence or absence of IGF2 indicated no toxicity under our experimental conditions (Supplementary Fig. S4L). We then determined whether polyQ_79_-EGFP aggregates were also secreted upon IGF2 treatment. Although the signal was low, we detected the presence of large polyQ_79_-EGFP aggregates outside the cell at basal levels using filter trap, a phenomenon that was incremented 2-fold in cells expressing IGF2 (Fig. 6B). However, we cannot discard that these aggregated form of polyQ_79_-EGFP were generated in the cell culture media after secretion or during the concentration process of the supernatants.

**Figure 6.**
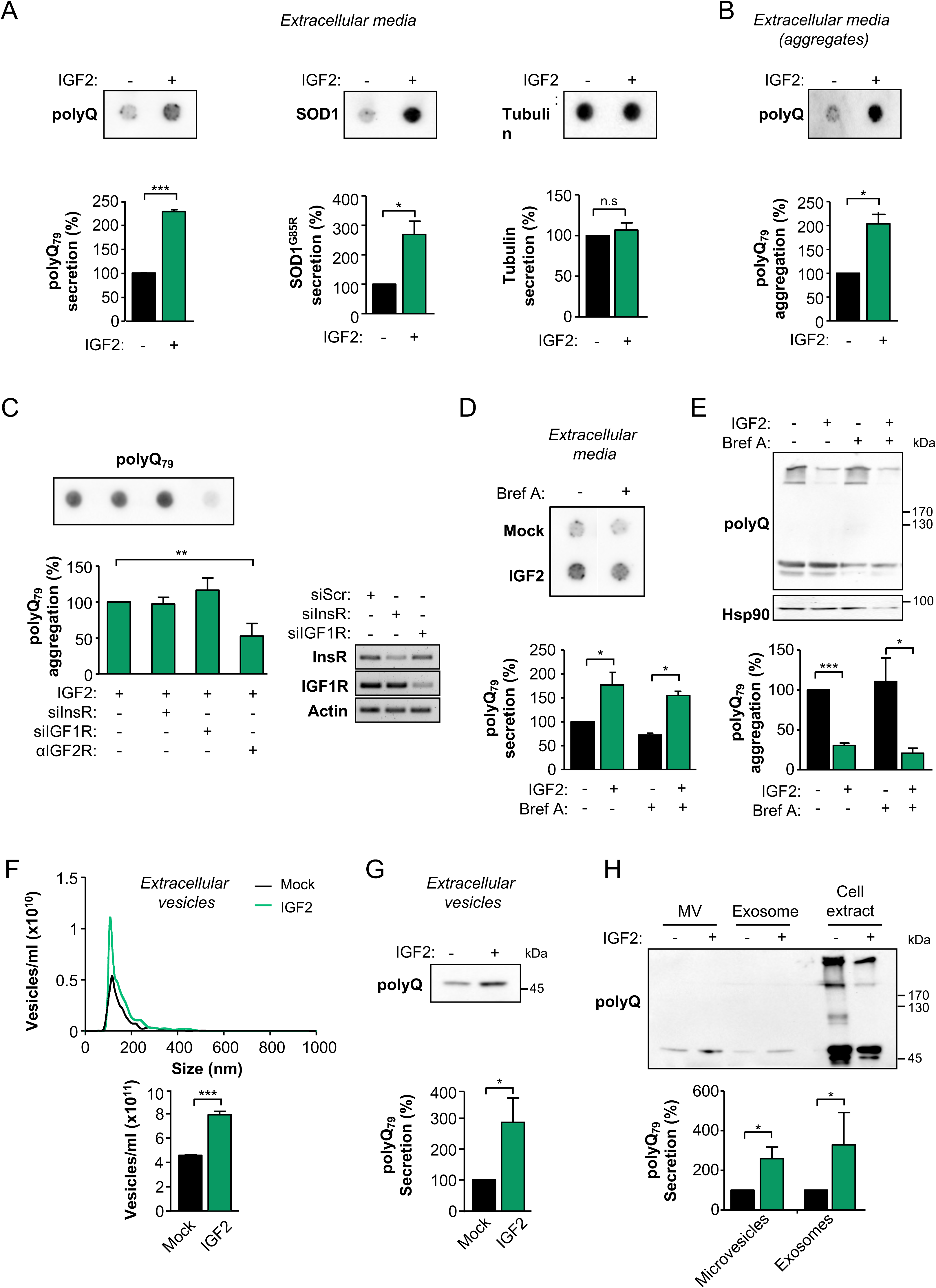
IGF2 enhances the extracellular release of polyQ_79_ and mHttQ_85_ through unconventional secretion. **(A)** Neuro2a cells were co-transfected with polyQ_79_-EGFP or SOD1_G85R_ plasmid and IGF2 plasmid or empty vector (Mock). After 16 h, cell culture media was replaced for Optimem and then collected after 24 h for dot blot or **(B)** filter trap analysis. **(C)** Neuro2a cells were transfected with siRNAs against InsR or IGF1R mRNAs. After 24 h, cells were co-transfected with polyQ_79_-EGFP and IGF2 expression vector or empty vector (Mock). Then, cell culture media was replaced for Optimem for 24 h. In addition, an anti-IGF2R was added when indicated. The presence of polyQ_79_-EGFP in the cell culture media analyzed by dot blot using an anti-GFP antibody and quantified. The knockdown of indicated proteins was confirmed in cell extracts using semi-quantitative PCR (right panel). **(D)** Neuro2a cells were co-transfected with polyQ_79_-EGFP and IGF2 plasmid or empty vector (Mock). Then, cell culture media was replaced for Optimem for 8 h and treated with 2 μM brefeldin A (Bref A) or vehicle for additional 16 h. Cell culture media was collected and analyzed by dot blot using an anti-GFP antibody (upper panel) and quantified (lower panel). Image was cropped from the same membrane and film exposure. **(E)** Neuro2a cells were co-transfected with polyQ_79_-EGFP plasmid and IGF2 plasmid or empty vector (Mock) for 8 h and then treated with 2 μM brefeldin A (Bref A) or vehicle for additional 16 h. PolyQ_79_-EGFP aggregates were analyzed in cell lysates by western blot using anti-GFP antibody. Hsp90 expression was monitored as loading control. PolyQ_79_-EGFP aggregates were quantified and normalized to Hsp90 (lower panel). **(F)** Extracellular vesicles from Neuro2a were analyzed by NanoSight nanotracking analysis and plotted by size (upper panel) and total concentration (lower panel) in cells expressing or not with IGF2. **(G)** Extracellular vesicles from Neuro2a cells co-transfected with polyQ_79_-EGFP and IGF2 expression vectors or empty vector (Mock) were concentrated followed by western blot analysis. **(H)** Isolated microvesicles (MV) and exosomes, together with total cell extracts from Neuro2a cells co-transfected with polyQ_79_-EGFP and IGF2 expression vectors or empty vector (Mock) were analyzed by western blot. In all quantifications, average and SEM of at least three independent experiments are shown. Statistically significant differences detected by two-tailed unpaired *t*-test (***: *p* < 0.001; **: *p* < 0.01; *: *p* < 0.05).

Given that IGF2 is a soluble factor, we evaluated which receptors mediated the enhancement of polyQ secretion. Thus, we used siRNAs to interfere insulin and IGF1 receptors (InsR and IGF1R), in addition to a blocking antibody to target the IGF2 receptor (IGF2R). Remarkably, antagonizing IGF2R decreased almost in half the levels of polyQ_79_-EGFP secretion, whereas the knockdown of InsR or IGF1R did not have any effect (Fig. 6C). Taken together, these results indicate that IGF2 signaling reduces the load of polyQ_79_-EGFP in the cell through the engagement of IGF2R, resulting in the extracellular disposal of polyQ proteins.

While the precise mechanisms of mHtt secretion are not completely understood, several possibilities have been suggested, including synaptic vesicle release (*41*), vesicular transport (*42*), exosomes/extracellular vesicles release (*43, 44*), exophers (*45*) and secretory lysosomes (*40*). Since the initial link connecting IGF2 with mHtt was through the UPR, we evaluated if the polyQ peptide is secreted by the conventional secretory pathway. Blocking ER to Golgi trafficking with brefeldin A was unable to ablate polyQ_79_-EGFP secretion in IGF2 stimulated cells (Fig. 6D, E). We then explored the presence of polyQ_79_-EGFP in extracellular vesicles. First, we used Nanoparticle Tracking Analysis system to determine the size distribution and concentration of vesicles in the cell culture media of cells expressing IGF2. We found enhanced release of vesicles in IGF2-expressing cells, showing an average diameter between 30-150 nm, whereas a minor portion (<1%) had a larger diameter (Fig. 6F). Interestingly, we observed that the population of vesicles increased under IGF2 expression picked around 100 nm, suggesting an enhancement of exosome release (*46*). Thus, we purified extracellular vesicles followed by western blot analysis and observed higher content of polyQ_79_-EGFP in vesicles obtained from IGF2-expressing cells (Fig. 6G).

We then isolated microvesicles and exosomes-enriched fractions through sequential centrifugation. Remarkably, increased levels of polyQ_79_-EGFP were detected in both fractions in cells expressing IGF2 using western blot analysis (Fig. 6H, see controls of purification in Supplementary Fig. S4M, N). Importantly, mostly monomeric and soluble forms of polyQ_79_-EGFP were secreted through microvesicles and exosomes, and no detergent insoluble aggregates were observed in this analysis. Thus, IGF2 signaling enhances the secretion of polyQ_79_-EGFP through exosomes and microvesicles, reducing the levels of polyQ_79_-EGFP inside the cell.

### Cytoskeleton remodeling contributes to polyQ secretion induced by IGF2

To define possible downstream effectors mediating the effects of IGF2 signaling in abnormal protein aggregation, we employed an unbiased approach to screen for global changes in protein expression. To characterize proteome remodeling upon stimulation with IGF2, we applied quantitative proteomics using Tandem Mass Tag-Multi-Dimensional Protein Identification Technology (MuDPIT) (*47*). Using this approach, we identified a total of 199 proteins that changed their expression levels with a *p* value < 0.05 and a minimum log2 > 0.1-fold change (Fig. 7A and Supplementary Table S2). IGF2 expression did not result in major proteomic changes. However, functional enrichment analysis revealed that IGF2 expression triggered fluctuations in proteins related to macromolecular complex disassembly, protein transport and cytoskeleton reorganization (Fig. 7B). A cluster of proteins modified by IGF2 were related to actin cytoskeleton dynamics and regulation, including several actin binding proteins, Rho GTPases, vimentin, dynactin, dyneins, among other factors (Supplementary Table S3).

**Figure 7.**
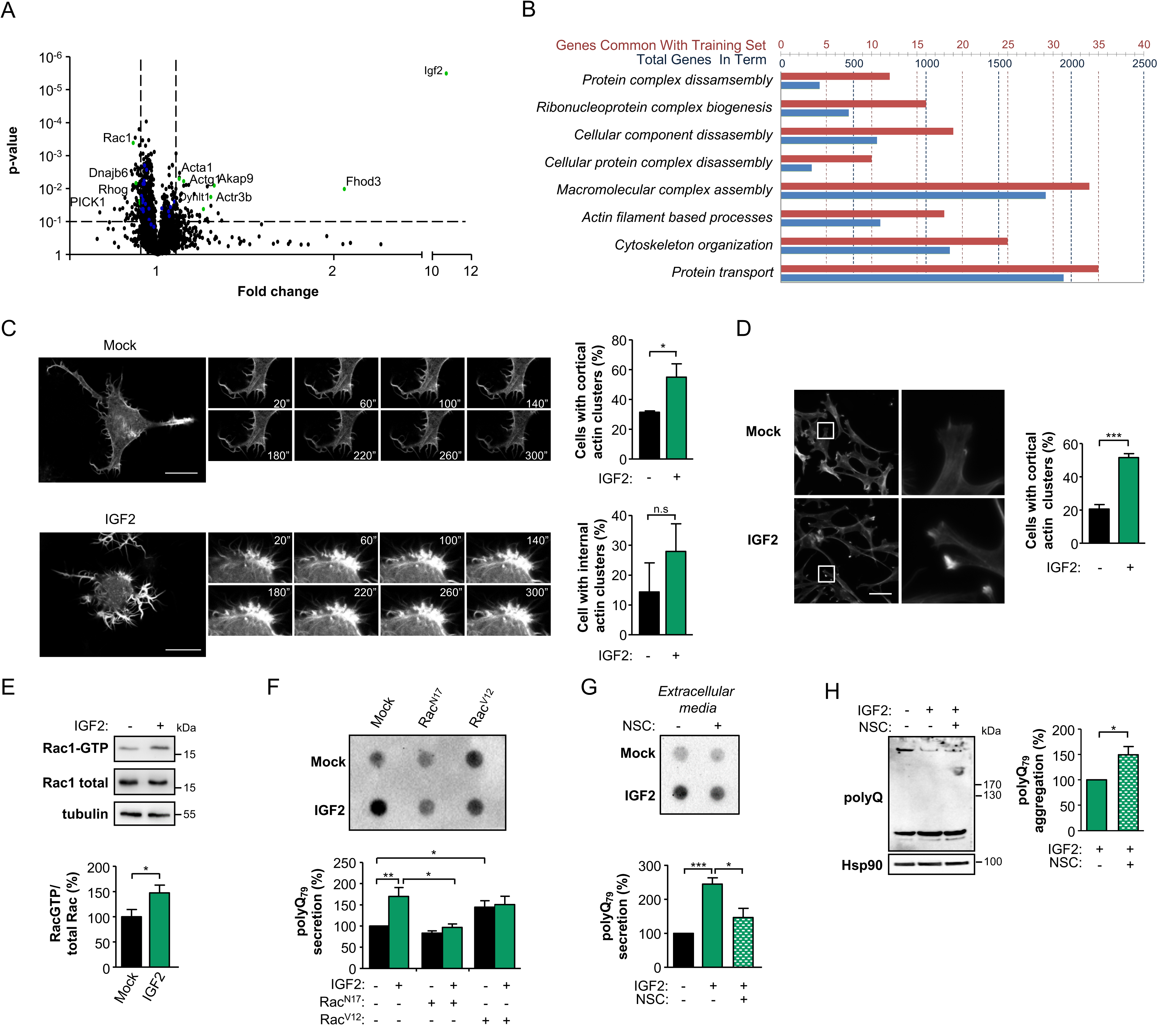
IGF2 signaling induces the release of polyQ_79_ aggregates through cytoskeleton remodeling. **(A)** Quantitative proteomics was performed in protein extracts derived from Neuro2a cells transiently transfected with IGF2 or empty vector (Mock) for 24 h. Data was analyzed and plotted in a volcano graph as fold-change. Vertical dashed lines indicate a log2 > 0.1 of fold change. Proteins related to actin cytoskeleton function are highlighted in green. Dots in blue represent associated proteins to actin cytoskeleton with a log2 < 0.1 fold-change. **(B)** Bioinformatics analysis of quantitative proteomics to uncover molecular processes affected by IGF2 expression after a functional enrichment analysis. Blue scale refers to the number of proteins that are known to be involved in each pathway. Red scale refers to the number of proteins altered in IGF2 overexpressing cells for each pathway. **(C)** Neuro2a cells were transfected with a plasmid encoding Life-Actin to monitor actin cytoskeleton dynamics. Cells were plated onto fibronectin-coated plates and recorded by time-lapse confocal microscopy every 40 s for 5 min. Time-lapse microscopy was performed after treatment with IGF2-enriched media (left panel). Quantification of cortical and internal actin clusters is shown (right panel). **(D)** Phalloidin-rhodamine staining of MEF cells after 5 min of treatment with IGF2-enriched media. Scale bar, 50 µm (right panel). Quantification of actin clusters is presented (left panel). **(E)** Neuro2a cells were treated with IGF2-enriched media or control media derived from Mock transfected cells. After 5 min, cells were lysed and cell extracts prepared to measure Rac1-GTP levels by pull-down assay using GST-CRIB domain. Pull down and total cell lysates were evaluated by western blot using anti-Rac1 antibody. Total Rac1 and tubulin monitored as loading control (upper panel). Rac1-GTP levels were quantified and normalized to total Rac1 and tubulin (lower panel). **(F)** Neuro2a cells were transfected with Rac_N17_ or Rac_V12_ or empty vector (Mock). 24 h later cells were co-transfected with expression vectors for polyQ_79_-EGFP and IGF2 or empty vector (Mock) and then incubated with Optimem. The presence of polyQ_79_-EGFP in the cell culture media was determined using dot blot (upper panel) and quantified (lower panel). **(G)** Neuro2a cells were co-transfected with expression vectors for polyQ_79_-EGFP and IGF2 or empty vector (Mock). Then, cell culture media was replaced for Optimem for 8 h and treated with 100 μM NSC23766 (NSC) for additional 16 h. Cell culture media was collected and analyzed by dot blot using an anti-GFP antibody (upper panel) and quantified (lower panel). Image was cropped from the same membrane and film exposure. **(H)** Neuro2a cells were co-transfected with polyQ_79_-EGFP plasmid and IGF2 plasmid or empty vector (Mock) for 8 h and then treated with 100 μM NSC23766 (NSC) or vehicle for additional 16 h. PolyQ_79_-EGFP aggregates were analyzed in cell lysates by western blot using anti-GFP antibody. Hsp90 expression was monitored as loading control (left panel). PolyQ_79_-EGFP aggregates were quantified and normalized to Hsp90 levels (right panel). In all quantifications, average and SEM of at least three independent experiments are shown. Statistically significant differences detected by two-tailed unpaired *t*-test (***: *p* < 0.001; *: *p* < 0.05).

Since actin cytoskeleton is key to modulate protein secretion and vesicular trafficking (*48*), we decided to explore if IGF2 had any impact in the regulation of actin cytoskeleton dynamics using Lifeact, a fluorescent peptide designed to visualize polymerized actin in living cells (*49*). Neuro2a cells were plated on coverslips and recorded using time-lapse confocal microscopy. Unexpectedly, IGF2 treatment generated very fast changes in actin dynamics and cell morphology within minutes, reflected in increased development of filopodia and the appearance of actin clusters in the cytosol of the cell (Fig. 7C, see Supplementary videos 1 and 2). These results were confirmed in murine embryonic fibroblasts by visualizing the actin cytoskeleton in fixed cells stained with phalloidin-rhodamine (Fig. 7D).

Actin cytoskeleton dynamics are dependent on the activity of small GTPases from the Rho family (*50*). Among the small GTPases of the Rho family, Cdc42, Rac, and Rho are recognized as the most important regulators of actin assembly, controlling the formation of filopodia, lamellipodia and stress fibers (*51*). Considering the central role of Rac1 in cytoskeleton dynamics, we evaluated Rac activity upon IGF2 stimulation. Neuro2a cells were treated for 5 minutes with IGF2-enriched media and Rac1 activity was measured using pull-down assays using p21-activated kinase that binds specifically to the Rac1-GTP but not to the inactive form of Rac1 (Rac1-GDP) (*52*), followed by western blot analysis. These results demonstrate that the amount of active Rac1 coupled to GTP was increased very quickly after IGF2 treatment (Fig. 7E).

We tested the functional contribution of actin dynamics to the release of polyQ_79_-EGFP peptide into the extracellular space. Neuro2a cells were transiently transfected with both a dominant negative (Rac1_17N_) and a constitutive active (Rac1_V12_) forms of Rac1. Remarkably, Rac1_17N_ was able to block IGF2-enhanced secretion of polyQ_79_-EGFP (Fig. 7F). In contrast, Rac1_V12_ expression facilitated polyQ_79_-EGFP secretion at basal levels, whereas no differences were observed in IGF2-stimulated cells suggesting that the pathway was already saturated (Fig. 7F). In agreement with these findings, inhibition of Rac1-GTPase activity with NSC23766 significantly decreased the secretion of polyQ_79_-EGFP (Fig. 7G). These effects paralleled with an increase in the levels of intracellular polyQ_79_-EGFP aggregation in cells treated with IGF2 in the presence of NSC23766 (Fig. 7H). Taken together, these results suggest that IGF2 signaling triggers rapid changes in cytoskeleton dynamics that result in the routing of polyQ_79_-EGFP peptide into the extracellular space.

### IGF2 reduces mHtt aggregation in vivo

Based on the significant reduction of mHtt aggregation observed in cells overexpressing IGF2, we moved forward to develop a therapeutic strategy to deliver IGF2 into the brain of HD mice using the stereotaxic injection of AAVs to then assess the impact on mHtt levels. We previously developed an animal model of HD to monitor mHtt aggregation based on the local delivery into the striatum of a large fragment of mHtt of 588 amino acids containing 95 glutamine repetitions fused to monomeric RFP (Htt588Q_95_-RFP) (*53*). To validate the effects of IGF2 in the aggregation of this mHtt construct, we first performed co-expression experiments in Neuro2a cells followed by fluorescent microscopy and western blot analysis. IGF2 expression decreased mHtt aggregation in both experimental settings (Supplementary Fig. S5A).

To determine the possible impact of IGF2 in mHtt aggregation levels *in vivo*, we performed unilateral stereotaxic injections of a mixture of AAVs to deliver Htt588Q_95_-RFP together with IGF2 or empty vector (Mock) into the striatum (Figure 8A, left panel). Two weeks after AAV delivery, mice were euthanized, and the striatum was dissected for biochemical analysis. We confirmed the overexpression of IGF2 in the striatum using PCR (Supplementary Fig. S5B). Local expression of IGF2 in adult mice resulted on a marked decrease of mHtt aggregation of near 90% when evaluated by western blot analysis (Fig. 8A, middle and right panel). Consistent with our cell culture experiments, monomeric forms of mHtt were also reduced in the brain of HD mice injected with AAV-IGF2 (Fig. 8A, middle and right panel). To assess the impact of overexpressing IGF2 on neuronal survival, we measured DARPP-32 levels as a marker of medium spiny neuron viability (*22*). Interestingly, significantly higher levels of DARPP-32 were detected in AAV-IGF2-treated animals when compared to control virus (Fig. 8a, middle and right panel).

**Figure 8.**
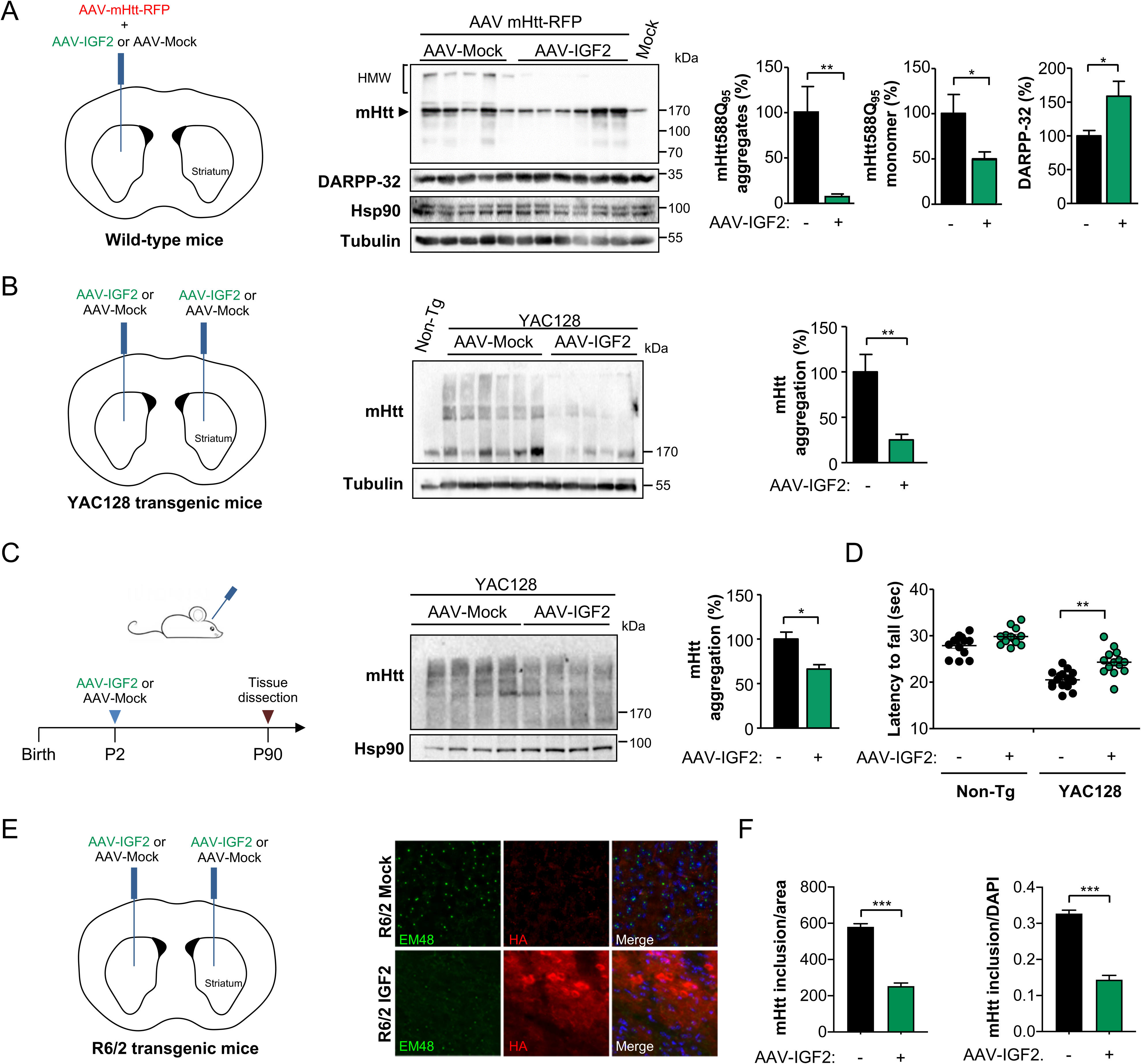
Gene therapy to deliver IGF2 into the striatum reduces mHtt aggregation in HD mouse models. **(A)** Three-month-old wild type mice were co-injected with adeno-associated viral (AAV) vectors expressing mHtt588Q_95_-RFP and IGF2 or empty vector (Mock). Two weeks later, the striatum was dissected and mHtt aggregation was analyzed in total protein extracts by western blot analysis using anti-polyQ antibody. DARPP32 levels were also determined in the same samples. Hsp90 and tubulin expression were monitored as loading control. The presence of the monomer is indicated with an arrowhead. mHtt588Q_95_-RFP aggregates and monomers or DARPP32 levels were quantified and normalized to Hsp90 levels (lower panel) (AAV-Mock: n = 5; AAV-IGF2: n = 6). **(B)** Three-month-old YAC128 transgenic mice were injected with AAV-IGF2 or AAV-Mock into the striatum using bilateral stereotaxis. Four weeks later, the striatum was dissected and mHtt aggregation monitored by western blot using anti-polyQ antibody. Tubulin expression was monitored as loading control (upper panel). mHtt aggregates levels were quantified and normalized to tubulin levels (lower panel) (AAV-Mock: n = 6; AAV-IGF2: n = 5). **(C)** YAC128 mice were injected at postnatal stage (P1-P2) into the ventricle with AAV-IGF2 or AAV-Mock. Three months later, striatum was dissected and mHtt aggregation levels were analysed by western blot using anti-polyQ antibody. Hsp90 levels were determined as loading control (left panel). mHtt aggregates levels were quantified and normalized to Hsp90 levels (right panel) (AAV-Mock: n = 5; AAV-IGF2: n = 4). **(D)** YAC128 and littermate control mice were injected with AAVs as indicated in (c). Motor performance was monitored once every two weeks from 3 months to 8 months old. The analysis shows the average of the group at each time point (AAV-Mock: n = 11; AAV-IGF2: n = 9). Statistically significant differences detected by two-tailed unpaired *t*-test (**: *p* < 0.001; *: *p* < 0.05). **(E)** R6/2 mice were injected at 4 weeks of age with AAV-IGF2 of AAV-Mock into the striatum using bilateral stereotaxis. Four weeks after the injection, mice were perfused, the brain was extracted and coronal slices from the striatum were obtained. Mutant huntingtin and HA-epitope were detected using the EM48 and anti-HA antibodies, respectively. Nuclei were stained using DAPI. High-resolution images of the slices were obtained using fluorescence microscopy and quantification of mutant huntingtin and DAPI-positive cells was performed using Image J software. **(F)** Quantification of the number of mutant huntingtin inclusions was performed by total area (left panel) and number of DAPI-positive cells (right panel). (AAV-Mock: n = 4; AAV-IGF2: n = 4). Statistically significant differences detected by two-tailed unpaired t-test (***: *p*<0.001; **: *p*<0.01; *: *p*<0.05)

Considering our positive results obtained using the viral HD model, we then evaluated our gene therapy strategy to deliver IGF2 in the YAC128 transgenic model on a heterozygous condition. This mouse model expresses full-length mHtt with 128 tandem glutamines using an artificial chromosome that contains al endogenous regulatory elements (*54*). First, we performed bilateral stereotaxic injections of AAV-IGF2 or empty vector into the striatum of 3-month old YAC128 mice followed by biochemical analysis of brain extracts four weeks later (Fig. 8B, left panel). This strategy led to a significant reduction of full-length mHtt expression in the striatum, reaching near 80% decrease on average (Fig. 8B, middle and right panel). Because stereotaxic injections only transduce a restricted area of the striatum restricting the analysis on motor function, we employed a second route of AAV delivery to generate a global spreading of the viral particles through the brain to assess the impact of IGF2 on mHtt levels and motor control. The delivery of AAVs into the ventricle of new-born pups has been reported to result on an efficient spreading of the virus throughout the nervous system (*55*). We injected AAV-IGF2 or control vector in postnatal day 1 or 2 (P1-P2) YAC128 mice and dissected the striatum 3 months after (Figure 8C, left panel). Western blot analysis demonstrated that IGF2 expression significantly reduces the total levels of mHtt in the brain of YAC128 animals (Fig. 8C, middle and right panel; see controls for IGF2 expression in Supplementary Fig. S5C). Finally, we determined the functional consequences of administrating our IGF2-based gene therapy on the clinical progression of experimental HD. YAC128 mice develop a variety of functional deficits, including motor learning deficits on rotarod beginning at 2 months (*56*) and motor impairment starting at 4 months of age (*57, 58*). We monitored the motor performance of postnatal-treated YAC128 mice with control or AAV-IGF2 using the rotarod assay. We reported that this model shows a sustained impairment of motor control when assessed using the rotarod assay, with constant values for several weeks (*22*). Remarkably, delivery of IGF2 into the nervous system improved the average motor performance over time of HD transgenic mice (Fig. 8D).

To complement our *in vivo* validation, we took advantage of a third model, the R6/2 mice, a transgenic HD model that expresses exon 1 of human huntingtin containing ∼150 CAG repeats (*59*), which allows the visualization of intracellular mHtt inclusions. Thus, we evaluated the effects of IGF2 administration in the brain of R6/2 mice by performing bilateral stereotaxic injections of AAV-IGF2 or AAV-Mock into the striatum of 4 weeks old R6/2 mice, followed by tissue immunofluorescence analysis of the brains four weeks later (Figure 8E, upper panel). A strong reduction in the content of mHtt-positive inclusions was observed upon AAV-IGF2 administration (Figure 8E, lower panel). Quantification of these experiments indicated a significant reduction of near 60% in the brain of R6/2 mice treated with AAV-IGF2 (Fig. 8F). Taken together, these results indicate that the artificial enforcement of IGF2 expression in the brain reduces abnormal protein aggregation in different HD models.

### IGF2 levels are reduced in the striatum and blood samples of HD patients

Due to the dramatic effects of IGF2 on intracellular mHtt levels, we moved forward to explore the possible alterations on IGF2 expression in HD patient samples. We monitored the levels of IGF2 in human caudate-putamen derived from HD patients and age-matched control subjects obtained from the Harvard Brain Bank (Supplementary Table S4). Western blot analysis of protein extracts revealed a near 66% reduction on average of IGF2 levels in the brain of HD patients when compared to the control group (Fig. 9A).

**Figure 9.**
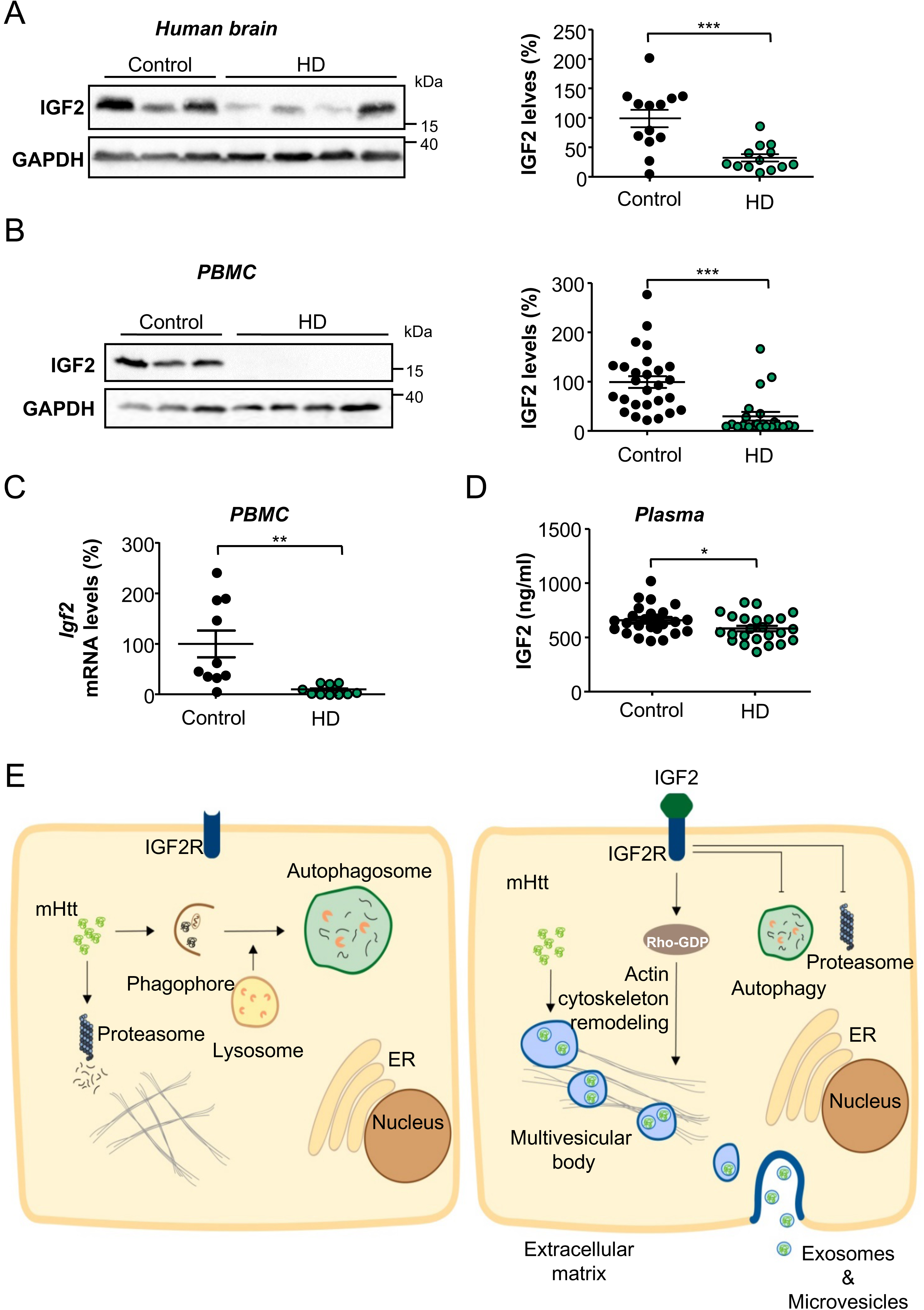
IGF2 levels are downregulated in the brain and blood of HD patients. **(A)** IGF2 protein levels were measured by western blot in human post mortem samples containing the caudate-putamen region from HD patients in stages 3 and 4. GADPH was monitored as loading control (left panel). IGF2 levels were quantified and normalized to GADPH levels (right panel). **(B)** Total protein extracts were generated from freshly isolated peripheral blood mononuclear cells (PBMCs) from HD patients and control subjects. IGF2 expression levels were determined using western blot analysis. GADPH was determined as loading control (left panel). IGF2 levels were quantified and normalized to GADPH levels (right panel). **(C)** *Igf2* mRNA levels were measured by real-time PCR in PBMCs obtained from HD patients and control subjects. **(D)** IGF2 content in plasma from blood obtained from HD patients and control subjects was measured by ELISA. In all quantifications, statistically significant differences detected by two-tailed unpaired *t*-test (***: *p* <0.001; **: *p* < 0.01; *: *p* < 0.05). **(E)** Proposed model: IGF2 reduces mHtt levels through unconventional secretion. In HD patients, the downregulation of IGF2 expression enhances the content of mHtt (left panel). When IGF2 is upregulated using gene therapy, IGF2R signalling enhances the secretion of mHtt into the extracellular space involving actin cytoskeleton remodelling and possibly microvesicles and exosomes (right panel).

Considering the unexpected decrease of IGF2 levels in HD postmortem brain tissue, we evaluated the presence of IGF2 in peripheral blood mononuclear cells (PBMCs) from HD patients. We obtained blood samples from a cohort of 24 patients recruited in the Enroll-HD international platform (Supplementary Table S5). Although control PBMCs presented a clear expression of IGF2, HD-derived cells showed reduced IGF2 levels, observing an almost 80% decrease on its protein levels using western blot analysis (Fig. 9B). These results were confirmed when mRNA levels of IGF2 were measured in the same samples, observing a near 90% decreased in its levels (Fig. 9C). Since IGF2 is a soluble secreted factor and its plasma levels have been suggested as a possible biomarker of cancer (*60, 61*), we decided to measure the quantity of IGF2 in plasma from HD patients using ELISA. This analysis revealed a slight but significant decrease in the amount of circulating IGF2 present in the plasma samples derived from HD patients when compared to control subjects (Fig. 9D), which may be related to the different contribution of tissues and cell types to plasmatic IGF2 levels. Taken together, these results suggest that IGF2 levels are drastically reduced in the brain and blood cells of HD patients.

## Discussion

Proteostasis impairment is observed in a variety of neurodegenerative diseases and is a central hallmark of aging (*62*). Our previous studies uncovered an unexpected connection between the UPR and autophagy, two central nodes of the proteostasis network, where a dynamic balance between both pathways sustains cellular function (*18, 22*). However, it remained to be defined if other mechanisms may underlie the beneficial targeting of the UPR in neurons. To identify novel disease modifiers, we performed a gene expression profile analysis of striatal and cortical areas of XBP1 ablated animals and uncovered the upregulation of *Igf2* as the major hit. This prompted us to investigate the consequences of manipulating IGF2 in the context of PMDs using HD as a proof-of-concept. Based in our findings, and other studies, we speculate that growth factor signaling may play an important role in boosting relevant pro-survival effectors that fine tune the proteostasis network (*63*).

IGF1 and IGF2 are mitogenic polypeptides with structural homology to insulin. IGF2 binds with higher affinity to IGF2R, but can also associate with lower affinity to IGF1R and the insulin receptor (*64, 65*). IGF2 regulates cell proliferation, growth, differentiation and survival (*66*). In general, the biological effects of IGF2 have been historically mapped to the IGF1R, and to a lower extent to the insulin receptor, impacting cell survival and proliferation (*67*). Unlike insulin and IGF1 receptors, IGF2R does not have intrinsic tyrosine kinase activity and has higher affinity for IGF2 than IGF1, and does not bind insulin (*68*). IGF2R controls extracellular IGF2 levels as it promotes its endocytosis towards lysosomal-mediated degradation (*68*). However, IGF2R can initiate specific responses affecting various cellular processes, possibly involving the activation of heteromeric G proteins and downstream calcium signaling, in addition to the activation of PKC and MAP kinases (reviewed in (*68*)). Intracellular IGF2R also controls the uptake and trafficking of lysosomal enzymes from the trans-Golgi network to the lysosomes (*68*). While fetal IGF2 is abundant, its levels are decreased after birth and are also further reduced during aging (*29*). In humans, altered dosage of the *IGF2* gene can result in developmental problems and its deregulation has been widely correlated with cancer (*69*). In adults, *IGF2* is almost exclusively expressed in the brain, especially in the choroid plexus, the brain vasculature and meninges (*70*). Initial assessment of the significance of *IGF2* expression to animal physiology was provided by the observation that full knockout animals develop growth defects (*71*), whereas it overexpression generates embryonic lethality (*72, 73*).

Recent findings highlight the physiological function of IGF2 in cognition, neuronal differentiation and survival. Several studies have demonstrated the relevance of IGF2 to memory consolidation (*74–76*), in addition to memory extinction (*77*), cognitive and social learning (*78, 79*) and brain stem cells proliferation in adults related to learning, memory and anxiety (*80*). At the molecular level, IGF2 regulates the formation of dendritic spines (*29*) and synapses (*76*), in addition to control adult neurogenesis in the hippocampus (*81*). At the dentate gyrus, the autocrine action of IGF2 may explain the proliferative effects on neural stem cells (*81*). Importantly, the effects of IGF2 in learning, memory and neurogenesis have been mapped to the stimulation of IGF2R. Thus, it might be feasible that IGF2 contributes to neuroprotection in HD through additional mechanisms, improving synaptic plasticity, connectivity, neuronal survival and neuronal function. We are currently addressing this hypothesis by assessing the possible impact of IGF2 in the clinical progression of HD using gain- and loss-of-function. Our current study proposes that IGF2 has a relevant role in controlling cellular proteostasis by fine-tuning the load of misfolded proteins through a novel mechanism involving the extracellular disposal of these misfolded proteins through extracellular vesicles (Figure 9E). We speculate that the reduction in the levels of soluble non-aggregated forms of mHtt and polyQ may shift the equilibrium in the amyloid cascade, reducing the proportion of abnormal protein aggregates inside the cell.

Our results suggest that the upregulation of IGF2 in animals with perturbed UPR may serve as a backup mechanism to sustain neuronal function when the adaptive capacity of the proteostasis network is severely compromised. We are currently investigating the mechanisms explaining the upregulation of IGF2 in XBP1 deficient brains. IGF2 expression is controlled by four different promoters and the presence of several mRNA binding proteins (*82*), in addition to the regulation by genomic imprinting according to a parental origin (*71, 83, 84*). IGF2 is also modulated in various pathological context including brain injury (*85*), schizophrenia (*86*), Alzheimeŕs disease (*29, 87*), in addition to normal aging (*29, 88*). Recent studies also indicated that the administration of IGF2 through gene therapy or the infusion of recombinant proteins into the brain ameliorates AD pathogenesis, reducing synaptic dysfunction (*28, 29*). Similarly, treatment of SOD1 transgenic mice with AAVs encoding for IGF2 delayed disease progression, improving motor neuron survival (*30*). These findings are consistent with the pro-survival role of IGF2 in motor neurons (*89*), in addition to its ability to induce axonal sprouting (*90, 91*). All these reports highlight the neuroprotective potential of IGF2 in various diseases affecting the nervous system.

Although several mechanisms have been proposed to be involved in the secretion of mHtt (*41–44*), a recent report suggested that secretory lysosomes are the major route for mHtt release into the extracellular space (*40*). Our results suggest that IGF2 signaling leads to a re-routing of mHtt, mostly the soluble forms, toward the secretion in extracellular vesicles. As a limitation of our study, we cannot exclude the possibility that IGF2 administration will enhance the propagation of abnormal mHtt conformations because our experimental settings were not designed to study disease spreading throughout the brain. Based on our results, we propose that soluble species of mHtt are secreted upon stimulation with IGF2, a form of the protein that is expected to lack the capacity to seed aggregation and propagate the protein misfolding process. It is most likely that the reduced aggregation observed in our experiments is due to a restricted availability of the soluble mHtt substrate, shifting the balance toward the accumulation of non-pathogenic mHtt species and reducing the overall intracellular levels of the protein. In addition, microglia has been shown to express IGF2R and IGF1R (*92*), opening the possibility that IGF2 may impact other cell types affected in HD. In fact, based on the data available in AD models (*28, 29*), we speculate that IGF2 administration may also enhance the extracellular clearance of mHtt by glial cells or by the action of extracellular proteases. Besides, recent studies suggested that mHtt is cleared faster in astrocytes than in neurons (*93*).

Our findings in human HD-derived tissue suggest that the downregulation of IGF2 during the progression of the disease may enhance or accelerate the accumulation of abnormal mHtt species. In contrast, increased plasmatic levels of IGF1 in HD patients predict a worst prognosis, involving a faster and stronger cognitive decline (*94, 95*). These effects were not observed when insulin levels were monitored in longitudinal studies (*95*). Moreover, analysis of YAC128 mice also revealed an increase in IGF1 expression in the brain and blood of symptomatic animals (*96, 97*). Since IGF2 levels were dramatically downregulated in human brain tissue derived from HD patients, we speculate that the delivery of IGF2 into the brain using gene therapy may serve as a strategy to restore its normal levels in the brain of HD patients.

Overall, our study places IGF2 as an interesting disease modifier agent and as an interesting candidate molecule for the treatment of HD. Since IGF2 is a soluble factor, the development of gene transfer strategies to augment IGF2 levels in the brain may emerge as an attractive approach for future therapeutic development. Based on the available literature and our current study we speculate that IGF2 administration to HD patients may have important beneficial consequences beyond proteostasis control and mHtt aggregation, including multiple points of action such as improvement of neuronal connectivity and synaptic plasticity, in addition to enhance axonal regeneration, neuronal survival, neurogenesis/tissue repair and improvement in motor control.

## Materials and methods

### Animal care

XBP1_flox/flox_ mice were crossed with mice expressing Cre recombinase under the control of the Nestin promoter to achieve the conditional deletion of XBP1 in the nervous system (XBP1_cKO_) (*27*). We employed as HD models the R6/2 transgenic mice, expressing exon 1 of the human huntingtin gene carrying approximately 150 CAG repeat expansions (*98*), and the full-length human Htt transgenic mice with 128 CAG repetitions termed YAC128 (*54*), both obtained from The Jackson Laboratory. To generate experimental animals, XBP1_cKO_ mice were crossed with the YAC128 model on FVB background and every generation breed to pure background XBP1_cKO_ mice for four to six generations to obtain experimental animals. For proper comparison, all biochemical and behavioral analysis were performed on groups of littermates of the same breeding generation. Unless indicated, YAC128 and littermate control animals were used and maintained on a pure C57BL/6J background. Mice were maintained in a quiet, ventilated, and temperature-controlled room (23°C), with a standard 12 h light cycle, and monitored daily. Mice were housed in polystyrene solid bottom plastic cages fitted with a filtertop. Mice were fed with LabDiet pellets and drinking water ad libitum. For euthanasia, mice received CO_2_ narcosis. Animal care and experimental protocols were performed according to procedures approved by the “Guide for the Care and Use of Laboratory Animals” (Commission on Life Sciences. National Research Council. National Academy Press 1996), the Institutional Review Board’s Animal Care and Use Committee of the Harvard School of Public Health, the Bioethical Committee of the Faculty of Medicine, University of Chile (CBA 0670 FMUCH) and the Bioethical Committee of the Center for Integrative Biology, University Mayor (07–2017). We used XBP1_cKO_ (N = 17) and litter matte control (N = 12) animals to gene expression analysis. For gene therapy analysis, we used 4-6 YAC128 mice per experimental group and 8 R6/2 mice per experimental group.

### Microarray

XBP1_cko_ animals were euthanized at 12-14 months of age and perfused with heparinized saline buffer. Brains were removed and placed in RNAlater. Brain cortex and striatum were dissected and mouse whole-genome profiling was performed using the Illumina BeadChip® platform (Illumina; San Diego, CA). According to the manufacturer, probes on the Illumina MouseWG-6 v2.0 Expression BeadChip were derived from the National Center for Biotechnology Information Reference Sequence (NCBI RefSeq) database (Build 36, release 22), supplemented with probes derived from the Mouse Exonic Evidence Based Oligonucleotide (MEEBO) set, as well as exemplar protein-coding sequences described in the RIKEN FANTOM2 database. The MouseWG-6 v2.0 BeadChip contains the full set of the MouseRef-8 Expression BeadChip probes with an additional 11,603 probes from the above databases. Raw data was quantile normalized and differentially expressed genes identified using the ArrayAnalysis software (*99*). Genes that were significantly up or down regulated were identified using Significance Analysis of Microarrays (SAM) (*100*). SAM assigns a score to each gene based on a change in gene expression relative to the standard deviation of repeated measurements. For genes with scores greater than an adjustable threshold, SAM uses permutations of the repeated measurements to estimate the percentage of genes identified by chance the false discovery rate (FDR). Analysis parameters were set to 800 permutations, and genes with q-value =0% were used for genomics analysis.

To identify functional connections between deregulated transcripts, both network and pathway analyses of the genes filtered by microarray were performed using Ingenuity Pathways Analysis (IPA; QIAGEN Inc., https://www.qiagenbioinformatics.com/products/ingenuitypathway-analysis). The significance of networks was calculated by integrated Ingenuity algorithms. IPA computes a score for each gene network according to the fit of that network to the user-defined set of “focus genes.” The score is derived from a P value and indicates the likelihood of the focus genes in a network being found together due to random chance. Relevant pathways with p-values less than 0.05 were considered. In addition, IPA compares the direction of change for the differentially expressed genes with expectations based on the literature (cured in the Ingenuity Knowledge Base) to predict an integrated direction of change for each function, using the z-score algorithm. For heatmap representation of deregulated transcripts the software HeatmapGenerator was used (*101*).

### RNA isolation, RT-PCR and real-time PCR

Total RNA was prepared from tissues or cells placed in cold PBS using Trizol following the manufacturer’s instructions (Life Technologies). The cDNA was synthesized with SuperScript III reverse transcriptase (Life Technologies) using random primers p(dN)6 (Roche). Quantitative PCR reactions were performed using standard protocols using EvaGreen^TM^ in the Stratagene Mx3000P system (Agilent Technologies). The relative amounts of mRNAs were calculated from the values of comparative threshold cycle by using Actin as a control. PCR or RT-PCR were performed using the following primers: For mouse: *Igf2* 5’-GTCGCATGCTTGCCAAAGAG-3’ and 5’-GGTGGTAACACGATCAGGGG-3’, mouse *Actin* 5’-TACCACCATGTACCCAGGCA-3’ and 5’-CTCAGGAGGAG AATGATCTTGAT-3’, *Igf1* 5’-AAAGCAGCCCCGCTCTATCC-3’ and 5’-CTTCTGAGTCTTGGGCATGT A-3’, *Acot1* 5’-TGCAAAGCCCTCTGGTAGAC-3’ and 5’-CTCGCTCTTCCAGTTGTGGT-3’, *insulin receptor 5’-* TCAAGACCAGA CCGAAGATT-3’ and 5’-TCTCGAAGATAACCAGGGCATAG-3’, *Igf1r* 5’-CATGTGCTGGCAGTATAACCC-3’ and 5’-TCG GGA GGC TTG TTC TCC T-3’. For human *IGF2* 5’-GTGCTGTTTCCGCAGCTG-3’ and 5’-AGGGGTCGACACGTCCCTC-3’, *Actin* 5’-GCGAGAAGATGACCCAGATC-3’ and 5’-CCAGTGGTACGGCCAGAGG-3’, HA 5’-TAGACGTAATCTGGAACATCG-3’.

### Reagents, plasmids and transfection

Lactacystin, MG-132, chloroquine, 3-methyladenine, bafilomycin A_1_, brefeldin A and recombinant insulin were purchased from Sigma. NSC23766 was purchased from Santa Cruz. GW4869 was purchased from Cayman Chemicals. Phalloidin was purchased from Molecular Probes, Invitrogen. Blocking antibody for IGF2R was purchased from Santa Cruz (sc-25462). Cell media and antibiotics were obtained from Invitrogen (MD, USA). Fetal calf serum was obtained from Hyclone and Sigma. All transfections for plasmids were performed using the Effectene reagent (Qiagen). DNA was purified with Qiagen kits. PolyQ_79_-EGFP is a 79 polyglutamine tract in-frame fused to EGFP in the N-terminal; GFP-mHttQ_85_ contains the first 171 amino acids of the first exon of the huntingtin gene with a tract of 85 glutamines fused to GFP in the N-terminal, was kindly provided by Dr. Paulson Henry (University of Michigan). Myc-tagged full-length huntingtin with a 103 polyglutamine region (FL Htt103Q-myc) is in pcDNA3, and was kindly provided by the Coriell Institute. pAAV-mHttQ_85_-mRFP contains the first 588 amino acids of the huntingtin gene with a tract of 85 glutamines, fused to mRFP (Zuleta *et al.*, 2012). IGF2 cDNA was obtained from pSPORT6 kindly provided by Dr. Oliver Bracko and subcloned with or without the HA epitope into a pAAV vector. SOD1_G85R_-EGFP was described before (*102*). siRNA pool for IGFR1 and InsR were purchased from Origene and transfections were made with Lipofectamine® RNAiMAX transfection Reagent from Invitrogene.

### Microscopy, western blot and filter trap analysis

Neuro2a and HEK293T cells were obtained from ATCC and maintained in Dulbecco’s modified Eagles medium supplemented with 5% fetal bovine serum and Penicillin/Streptomycin (Gibco). 3 x 10^5^ cells were seeded in 6-well plate and maintained by indicated times in DMEM cell culture media supplemented with 5% bovine fetal serum and non-essential amino acids. Treatment with autophagy, proteasome or exocytosis inhibitors was generally performed for 16 h unless indicated.

We visualized the formation of intracellular polyQ_79_-EGFP, GFP-mHttQ_85_ and Htt588Q_95_-RFP inclusions in live cells after transient transfection using epifluorescent microscopy. We quantified the intracellular inclusion using automatized macros done in Image J software. Protein aggregation was evaluated by western blot in total cell extracts prepared in 1% Triton X-100 in PBS containing proteases and phosphatases inhibitors (Roche). Sample quantification was performed with the Pierce BCA Protein Assay Kit (Thermo Scientific).

For western blot analysis, cells were collected and homogenized in RIPA buffer (20 mM Tris pH 8.0, 150 mM NaCl, 0.1% SDS, 0.5% Triton X-100) containing protease and phosphatase inhibitors (Roche). After sonication, protein concentration was determined in all experiments by micro-BCA assay (Pierce), and 25-100 µg of total protein was loaded onto 8 to 15 % SDS-PAGE minigels (Bio-Rad Laboratories, Hercules, CA) prior transfer onto PVDF membranes. Membranes were blocked using PBS, 0.1% Tween-20 (PBST) containing 5% milk for 60 min at room temperature and then probed overnight with primary antibodies in PBS, 0.02% Tween-20 (PBST) containing 5% skimmed milk. The following primary antibodies and dilutions were used: anti-GFP 1:1000 (Santa Cruz, Cat. n° SC-9996), anti-SOD1 (Cell signaling, Cat. n° 2770), anti-polyQ 1:1000 (Sigma, Cat. n° P1874), anti-HSP90 1:2000 (Santa Cruz, Cat. n° SC-13119), anti-GAPDH 1:2000 (Santa Cruz, Cat. n° SC-365062), anti-HA 1:500 (Santa Cruz, Cat. n°SC-805), anti-IGF2 1:1000 (Abcam, Cat. n° Ab9574), anti-LC3 1:1000 (Cell Signaling, Cat. n° 2775S), anti-p62 1:1000 (Abcam, Cat. n° Ab56416), anti-polyUbiquitin 1:5000 (Santa Cruz, Cat. n° SC-8017), anti-DARPP32 1:1000 (Millipore, Cat. n° ab10518), anti-tubulin 1:2000 (Oncogene, Cat. n° CP06), Rac1 clone 23A8 1:1000 (Millipore, Cat. n° 05-389), for cofilin and p-cofilin we used homemade antibody at a 1:1000 dilution (Dr. James Bamburg). Bound antibodies were detected with peroxidase-coupled secondary antibodies incubated for 2 h at room temperature and the ECL system.

For western blot analysis using FL Htt103Q-myc, cells were grown in 100 mm dishes and cell lysis was done in 100 µl of 1% SDS in PBS for 5 min at 95°C. Samples were then diluted 1:5 with 2% Triton X100, pelleted in a microcentrifuge for 30 min at 4°C and 50 µl of the supernatant was loaded on SDS–PAGE. After transfer to a nitrocellulose membrane, blocking was done in 5% low-fat milk and 0.1% Tween-20 in PBS for 1 hr. Incubation with the primary antibody was overnight at 4°C. The following primary antibodies and dilutions were used: anti-calnexin 1:1000 (Sigma-Aldrich, Cat. n° C4731), anti-Htt 1:1000 (Cell Signaling, cat n°2773S) and anti-myc (custom produced from 9E10 hybridoma). After three washes in 0.1% Tween-20 in PBS, incubation with the appropriate secondary antibody was for 1 hr at room temperature. After washing, ECL was performed and the membrane was exposed and quantified in a Bio-Rad ChemiDocXRS Imaging System (Bio-Rad, Hercules, CA).

For western blot analysis using HD MSNs protein extracts, MSNs were harvested in M-PER Mammalian Protein Extraction Reagent (Pierce) with cOmplete Mini EDTA-free protease inhibitor (1 tablet/10 mL) (Roche). Whole-cell lysates were sonicated with a 5 second pulse followed by a 5 second rest (X 5 times) at 40% amplitude. Samples were centrifuged at 14,000 rpm at 4°C for 20 min, and supernatant was collected and stored at −20°C. Protein concentrations were estimated using the BCA assay (Pierce). Cell lysates were denatured under reducing conditions by boiling 10–20 μg total protein with 1 μL of 1M DTT and 4X LDS sample buffer (Invitrogen) at 95°C for 10 min. SDS-PAGE was performed using NuPage 4-12% Bis-Tris gels (Invitrogen) or 3-8% tris-acetate gels. Gels were run in 1X MOPS or 1X tris-acetate SDS running buffer containing 500 μL of antioxidant (Life technologies) for 90 min, and transferred to 0.45 μm nitrocellulose membrane using 1X NuPage transfer buffer at 20 V for 14 h. Membranes were blocked with 5% non-fat milk in TBS with 0.1% Tween 20 (TBS-T) for 1 hr at room temperature (RT) and incubated with primary antibody reconstituted in 5% non-fat milk overnight at 4°C. The following primary antibodies and dilutions were used: anti-Htt 1:500 (Millipore, MAB2166) and anti-vinculin 1:500 (Sigma-Aldrich, Cat. n°V9131). Secondary antibodies were incubated for 2 h at RT. Blots were washed in TBS-T for 10 min (3X), and developed using Pierce ECL (Thermo Scientific). Vinculin served as loading control. Densitometry analysis was performed using ImageQuant TL v2005.

For western blots using SCA3 iPSCs-derived neurons, neuronal cells were washed in HBSS and scraped, cells were immediately frozen in liquid N2 followed by lysis in RIPA buffer (50 mM Tris, 150 mM NaCl, 0.2% Triton X-100) containing 25 mM EDTA. For fractionation, lysates containing 1–2 µg/µl total protein dissolved in 50 mM Tris, 150 mM NaCl, 0.2% Triton X-100, 25 mM EDTA (RIPA buffer) were centrifuged at 22,000g for 30 min at 4°C. The pellet fractions were separated from supernatants (Triton X-100-soluble fraction) and homogenized by sonication in RIPA buffer containing 2% SDS (SDS fraction). β-mercaptoethanol (5%) was added in all the samples and subsequently incubated at 99°C for 5 min. Gels were loaded with 10-20 µg of the Triton X-100 fraction and 40 µl of the SDS fraction. Proteins were resolved by SDS-PAGE, transferred to nitrocellulose membrane and then processed for western blotting. Membranes were subsequently incubated with HRP-conjugated secondary antibodies (Amersham) at 1:7000 dilution. Visualization was performed with enhanced chemiluminescence. Samples were probed with primary antibodies: Ataxin-3 (Aviva Systems Biology; Cat#ARP50507_P050), βIII-tubulin (Abcam; Cat#ab7751), GAPDH (Fitzgerald; Cat#10R-G109A).

For filter trap assays, protein extracts were diluted into a final concentration of SDS 1% and were subjected to vacuum filtration through a 96-well dot blot apparatus (Bio-Rad Laboratories, Hercules, USA) containing a cellulose acetate membrane with pores with a 0.2 μm diameter (Whatman, GE Healthcare) as described in (Torres et al., 2015). Membranes were then blocked using PBS, 0.1% Tween-20 (PBST) containing 5% milk and incubated with primary antibody at 4°C overnight. Image quantification was done with the Image Lab software from BioRad.

### Pulse-chase experiments

HEK293T cells were seeded in 6 well plates and transiently transfected using polyethylenimine (PEI) with pcDNA-GFP-HttQ_43_ and mRFP/IGF2 constructs (in a ratio of 1:9). We used these constructs instead of the polyQ_79_-EGFP peptide because this experiment was standardized using GFP-HttQ_43_. For pulse labeling of cells (_35_S incorporation), after 24 h of transfection cells were washed twice with pre-warmed pulse labeling medium (methionine and cysteine free DMEM (Gibco) containing 10% dialyzed FBS (Thermo Scientific), Glutamax (Life technologies) and Penicillin/ Streptomycin (Gibco) and kept for 45 min in pre-warmed pulse labeling medium to deplete the intracellular pool of cysteine and methionine. 0.1 mCi/mL _35_S (EasyTag™ EXPRESS35S Protein Labeling Mix) were added in pre-warmed pulse labeling medium and pulsed for different time periods. Before harvesting the cells, the radioactive solution was removed and cells were washed twice with 1 mL PBS containing 15 μM cycloheximide. Cells were scraped in 250 μL immunoprecipitation lysis buffer (150 mM NaCl, 50 mM Tris-HCl pH 7.5, 0.5% NP-40, 0.5 mM MgCl_2_, 3% glycerol, 0.9 mM DTT) and protease inhibitor cocktail (complete Sigma-Aldrich) and lysed by repetitive passage (8 times) through a syringe with 26-gauge needle (0.45 mm). Samples were centrifuged (20.000 x g) and the supernatant was collected. 50 μL cell lysate was kept for input. 800 μL TNE buffer (50 mM Tris-HCl pH 8.0, 150 mM NaCl, 1 mM EDTA) was added to rest of the sample, and the GFP-HttQ_43_ was immunoprecipitated using GFP-trap (Chormotek, gtma-100). GFP-trap beads were added to each sample and incubated overnight at 4°C. Beads were collected on a magnetic rack and washed twice in immunoprecipitation wash buffer (0.2% NP40, 0.2% SDS, protease inhibitors), and twice in high salt buffer (TNE with 500 mM NaCl). Beads were boiled in sample buffer (4% SDS, 10% 2-mercaptoethanol, 20% glycerol, 0.004% bromophenol blue and 0.125 M tris HCl) and analyzed using SDS page and western blotting or autoradiography. To determine half-life, 0.1 mCi _35_S was added to each well and incubated for 90 min. After washing chase medium (DMEM containing 10% FCS, Glutamax, P/S, 1 mg/mL methionine and 1 mg/mL cysteine (Gibco)) was added to cells and incubated for different time points. Sample preparations and immunoprecipitation of GFP-HttQ_43_ was performed as described for the pulse labeling experiments.

### mHtt secretion and dot blot detection

PolyQ_79_-EGFP secretion was analyzed as described before (*40*). In brief, to collect secreted proteins, Neuro2a cells were transiently transfected for a total period of 32 h, incubated in Optimem for the last 24 hrs. Cell culture media was collected and centrifuged for 5 min at 5000 r.p.m to eliminate cell debris, and then applied to a 96-well dot blot apparatus (Bio-Rad Laboratories, Hercules, USA) on a PVDF membrane. Membranes were then blocked and proteins were detected following the western blot protocol.

For the detection of FL Htt103Q-myc using dot blot, HEK293 cells were grown on 60 mm plates in 3 mL medium and transfected with plasmids for the expression of FL Htt103Q-myc together with IGF2 or empty vector. Cells and medium were collected 24 h post transfection. After washing with PBS cells were lysed in 200 µl of 1% SDS in PBS with protease inhibitor cocktail. After boiling for 5 min, sample duplicates (50 µl for cell lysates and 100 µl for medium) were loaded on a nitrocellulose membrane (pre-wet with PBS) in a dot blotting apparatus. After 2 washes with PBS the membrane was transferred into 5% milk blocking solution. Proteins were detected following the western blot protocol.

### Vesicle isolation and nanoparticle tracking analysis

Extracellular vesicles were isolated from cell culture media. Briefly, Neuro2a cells were cultured for 24 h in Optimem. Collected medium was then depleted of cells and cellular debris at 2,000 x g for 10 min at 4°C. Extracellular vesicles were isolated by centrifugation of the collected supernatant at 110,000 x g for 70 min at 4°C. The resultant pellet was resuspended in PBS 1X with proteases and phosphatases inhibitors cocktail. Vesicles were analyzed for size and concentration using nanoparticle tracking analysis (NanoSight, NS300/ZetaSizer instrument). Microvesicles and exosomes were isolated from cell culture media using Neuro2a. First, cell culture media was depleted of cells and cellular debris at 2,000 x g for 10 min at 4°C. Then, microvesicles were pelleted by centrifugation of the collected supernatant at 11,000 x g for 34 min at 4°C. Finally, exosomes were isolated from the collected supernatant from microvesicles purification by centrifugation at 110,000 x g for 70 min at 4°C. The resultant pellet was then washed in PBS and centrifuged again for 70 min at 110,000 x g at 4 °C. The resultant pellet was resuspended in PBS 1X with proteases and phosphatases inhibitors cocktail.

### Quantitative proteomics

Neuro2a cells in 6-well plates were transiently transfected with IGF2 or empty expression vectors for 24 h. Lysates were prepared in RIPA buffer containing proteases and phosphatases inhibitors cocktail (Roche). After extracts sonication, protein concentration was determined with BCA (Thermo Fisher). For each sample, 20 μg of lysate was washed by chloroform/methanol precipitation. Samples for mass spectrometry analysis were prepared as described (Plate *et al.*, 2016). Air-dried pellets were resuspended in 1% RapiGest SF (Waters) and brought up in 100 mM HEPES (pH 8.0). Proteins were reduced with 5 mM Tris(2-carboxyethyl) phosphine hydrochloride (Thermo Fisher) for 30 min and alkylated with 10 mM iodoacetamide (Sigma Aldrich) for 30 min at ambient temperature and protected from light. Proteins were digested for 18 h at 37°C with 0.5 μg trypsin (Promega). After digestion, the peptides from each sample were reacted for 1 h with the appropriate TMT-NHS isobaric reagent (Thermo Fisher) in 40% (v/v) anhydrous acetonitrile and quenched with 0.4% ammonium bicarbonate for 1 hr. Samples with different TMT labels were pooled and acidified with 5% formic acid. Acetonitrile was evaporated on a SpeedVac and debris was removed by centrifugation for 30 min at 18,000 x *g*. MuDPIT microcolumns were prepared as described (*103*). LCMS/MS analysis was performed using a Q Exactive mass spectrometer equipped with an EASY nLC 1000 (Thermo Fisher). MuDPIT experiments were performed by 5 min sequential injections of 0, 10, 20, 30, …, 100% buffer C (500 mM ammonium acetate in buffer A) and a final step of 90% buffer C / 10% buffer B (20% water, 80% acetonitrile, 0.1% formic acid, v/v/v) and each step followed by a gradient from buffer A (95% water, 5% acetonitrile, 0.1% formic acid) to buffer B. Electrospray was performed directly from the analytical column by applying a voltage of 2.5 kV with an inlet capillary temperature of 275°C. Data-dependent acquisition of MS/MS spectra was performed with the following settings: eluted peptides were scanned from 400 to 1800 m/z with a resolution of 30,000 and the mass spectrometer in a data dependent acquisition mode. The top ten peaks for each full scan were fragmented by HCD using a normalized collision energy of 30%, a 100 ms activation time, a resolution of 7,500, and scanned from 100 to 1,800 m/z. Dynamic exclusion parameters were 1 repeat count, 30 ms repeat duration, 500 exclusion list size, 120 s exclusion duration, and exclusion width between 0.51 and 1.51. Peptide identification and protein quantification was performed using the Integrated Proteomics Pipeline Suite (IP2, Integrated Proteomics Applications, Inc., San Diego, CA) as described previously (*47*).

### Rac1 pull-down assay and actin dynamics

Neuro2a cells were cultured for 2 days in 100-mm-diameter plastic plates (10_7_ cells/plate) and then stimulated with IGF2 conditioned medium for 5 min at 37°C and 5% CO**_2_**. For Rac1 activation assays the expression and purification of the CRIB domain (amino acids 67–150) of p21-activated kinase (Pak1) were performed as follows: BL21 (DE3) *E. coli* strains carrying pGEX-GST-CRIB were grown overnight at 37°C in Luria broth media containing ampicillin. Cultures were diluted 1:100 and grown in fresh medium at 37°C to an OD_600_ of 0.7. Next, IPTG was added to a final concentration of 1 mM, the cultures were grown for an additional 2 h and then samples were collected and sonicated in lysis buffer A (50 mM Tris-HCl, pH 8.0; 1% Triton X-100; 1 mM EDTA; 150 mM NaCl; 25 mM NaF; 0.5 mM PMSF and protease inhibitor complex (Roche)). Cleared lysates were affinity purified with glutathione-Sepharose beads (Amersham). Loaded beads were washed ten times with lysis buffer B (lysis buffer A with 300 mM NaCl) at 4°C. GST fusion protein was quantified and visualized in SDS-polyacrylamide gels stained with Coomassie brilliant blue. Purified loaded beads were incubated for 70 min at 4°C with 1 mg of either Neuro2a lysates using fishing buffer (50 mM Tris-HCl, pH 7.5; 10% glycerol; 1% Triton X-100; 200 mM NaCl; 10 mM MgCl_2_; 25 mM NaF and protease inhibitor complex). The beads were washed three times with washing buffer (50 mM Tris-HCl, pH 7.5; 30 mM MgCl_2_; 40 mM NaCl) and then re-suspended in SDS-PAGE sample buffer. Bound Rac1-GTP was subjected to western blot analysis.

To measure actin dynamics, Neuro2a cells were seeded onto fibronectin-coated 25-mm coverslips, transfected with EGFP-Lifeact using Lipofectamine® 2000 Transfection Reagent and imaged in HBSS medium supplemented with HEPES using a confocal microscope (Zeiss LSM 710) with a 63×/1.4 NA oil-immersion objective at 37°C. Images were acquired every 30 s for 5 min, and the number of cells showing actin clusters were quantified. For phalloidin staining we used Phalloidin tetramethylrodamyne (Sigma) following the manufacturer’s instructions.

### Adeno-associated viral vector injections

All AAV (serotype 2) vectors were produced by triple transfection of HEK293T cells using a rep/cap plasmid and pHelper (Stratagene, La Jolla CA, USA), and purified by column affinity chromatography as previously described (*104*). For stereotactic injections, mice were anesthetized using a 100 mg/kg Ketamine, 10 mg/kg Xylazine mixture (Vetcom), and affixed to a mouse stereotaxic frame (David Kopf Instruments). Wild-type C57BL/6J male mice (3 months-old) were co-injected in the right striatum with 4 µl of virus AAV-Htt588Q_95_-mRFP and AAV-IGF2 or AAV-Mock in a relation 3:2 as we described before (*53*). The injection of AAVs suspension was performed at two points of the striatal region using a 5 µl Hamilton syringe (Hamilton) using the following coordinates: +0.7 mm anterior, +1.7 mm lateral and −3 to 3.2 mm depth, with a 1 µl/min infusion rate. After 2 weeks, mice were euthanized for biochemistry analysis. YAC128 mice (3 months-old) were injected bilaterally in a single point in the striatum with 2 µl of AAV-IGF2 or AAV-Mock. Stereotaxic coordinates with respect to the bregma used were: +0.7 mm anterior, +1.7 mm lateral and −3.1 mm depth, with a 1 µl/min infusion rate. After 1 month mice were euthanized for biochemistry analysis. Four weeks-old R6/2 mice were injected bilaterally in a single point in the striatum with 1.5 μl of AAV-IGF2 or AAV-Mock. Stereotaxis coordinates with respect to the bregma used were: +0.8 mm anterior, +1.8 mm lateral and 3.1 mm depth, with a 1 μl/min infusion rate. Four weeks after the injection, mice were euthanized for histochemical analysis.

### Rotarod test

Disease progression was monitored once every two weeks, using the rotarod assay as previously reported (Graham *et al.*, 2006). In brief, a training period was performed consisting of a first day of conditioning in the rotarod, following by three trials per day (with 2 h in between trials) over the course of 4 consecutive days. The data collected were considered an accurate reflection of the animal’s coordination. For each testing, mice were acclimated to the room for ∼15 min. The rotarod protocol consisted in (i) maximum speed, 40 r.p.m; (ii) time to reach maximum r.p.m, 120 s; (iii) time to ‘no fall’, 600 s; (iv) starting speed, 4 r.p.m; (v) trials per day, 3.

### Tissue preparation and analysis

For biochemistry of the striatum, mice were euthanized by CO_2_ narcosis, and brains were rapidly removed, rinsed with ice-cold 0.1 M phosphate buffer saline pH 7.4 (PBS), and placed on a cold surface to dissect different brain areas. Tissue samples were homogenized in 100 µl of ice-cold PBS (pH 7.4), supplemented with proteases and phosphateses inhibitor cocktail (Roche). The homogenate was divided for further mRNA and protein extraction following standard purification and quantification protocols. Protein extraction was performed in RIPA buffer (20 mM Tris pH 8.0, 150 mM NaCl, 0.1% SDS, 0.5% deoxycholate, and 0.5% Triton X-100) containing proteases and phosphatases inhibitor cocktail (Roche).

For tissue immunofluorescence analysis, mice were anesthetized and subjected to transcardial perfusion with ice-cold PBS (pH 7.4). Brains were removed and one hemisphere was fixed in 4% PFA for 24 h and then stored in a 30% sucrose solution. The other hemisphere was kept at −80 °C. Fixed hemispheres were cut using a cryostat (Leica CM1520) and 20 μm-thick coronal slices were obtained from the striatum. Free-floating slices were preserved in PBS-Sodium azide 0.02% at 4°C.

### Tissue immunofluorescence analysis

Brain slices were washed three times with PBS (pH 7.4) and then blocked at room temperature for 1 hr with blocking solution (BSA 0.5% and Triton 0.2% in PBS pH 7.4). Brain slices were incubated with EM48 1:1000 (Millipore, MAB5374) and HA 1:500 (Cell Signaling, cat n° C29F4) antibodies in blocking solution for 24 h at 4°C. Afterwards, slices were washed three times with PBS (pH 7.4) and then incubated for 2 h at room temperature with Alexa 488 goat anti mouse 1:1000 (Jackson ImmunoResearch, cat n° 115-545-166), Alexa 594 goat anti rabbit 1.1000 (Jackson ImmunoResearch, cat n° 111-545-144) and DAPI 1:10,000 (Merck, cat n° D9542). Finally, slices were washed three times with PBS (pH 7.4) and mounted in slides cover with gelatin using Fluoromount-G (Electron Microscopy Sciences, cat n° 17984-25).

### Microscopy and quantification of brain slices

High-resolution images (330.49 x 330.49 μm) from brain slices were obtained using an inverted Leica DMi8 light microscope using a 40X objective. For the quantification of mutant huntingtin inclusions and DAPI-positive cells, 20-30 z-stacks were acquired in three different regions of the striatum of a single slice for each experiment (six-nine slices per animal analyzed). Images were deconvolved using the microscope software and mutant huntingtin inclusions and DAPI-positive cells were quantified using automatized macros from the Image J software.

### Human samples

A total of 26 brain samples from the NIH NeuroBioBank were included in the study. Samples were obtained from 13 patients with clinical diagnosis of HD (grade 3 and 4 according to the neuronal loss observed) and 13 age-matched controls. Patients with HD met criteria proposed by the neuropathologist responsible in the Harvard Brain Tissue Resource Center (*105–107*). Samples of human adult peripheral blood (18 ml) were collected using BD vacutainer heparin tubes and processed in ≤4 hours after extraction. Plasma and PBMC were obtained by Ficoll gradient. Sampling was done according to the ethics committee form by the advisory board of the director of the local health service and the San Jose Hospital, University of Chile. Patients were recruited in the Enroll-HD project at the Movement disorder center (CETRAM) in Santiago, Chile (https://www.enroll-hd.org). Enroll-HD is a worldwide observational study for Huntington’s disease families. It will monitor how the disease appears and changes over time in different people and is open to people who either have HD or are at-risk. This platform collects standardized data, in the same way and using the same methods around the world. This project develops a comprehensive database of patients followed by a team of neurologists. Total, circulating IGF2 levels were measured in plasma from 49 individuals, using an ELISA IGF2 kit (CrystalChem, Cat. n° 80575).

### NSC culture and MSN differentiation

6-well plates were coated with 100 μg/ml poly-D-lysine (Sigma-Aldrich) followed by Matrigel (1:60) coating. HD and corrected C116 iPSC-derived neural stem cells (NSCs) were plated and cultured in Neural Proliferation Medium (NPM) (Neurobasal medium, 1X B-27 supplement (Life Technologies), 2mM L-Glutamine, 100U/ml penicillin, 100 μg/ml streptomycin, 10 ng/ml Leukemia inhibitory factor (LIF) (Peprotech, 300-05), 25 ng/ml basic Fibroblast Growth Factor (bFGF) (Peprotech, 100-18B), and 25 ng/ml Activin A (Peprotech, AF-120-14E) in humidified incubator under 37°C, 5% CO_2_. For differentiation into medium spiny neurons (MSNs), NSCs (when at ∼90% confluence) were treated with Synaptojuice A medium for 7 days followed by Synaptojuice B medium for 10 days at 37°C, 5% CO_2_ (*108*). 25 ng/ml Activin A was added to both Synaptojuice A and Synaptojuice B media. Half media change was performed every 2 days. NSCs were treated with AAV2-IGF2 or AAV2-PGK-empty (1 x 1011 VG per well) on day 1 of Synaptojuice A treatment. Untransduced MSNs were used as controls.

### Generation of iPSCs of SCA-3 patients with episomal vectors

Generation of iPSCs derived from patients with spinocerebellar ataxia class 3 was described previously (*109*). Human dermal fibroblasts (HDFs) were cultured in Dulbecco’s modified Eagle media (DMEM, Gibco) containing 10% fetal bovine serum (FBS), 1 mM non-essential amino acids (NEAAs), 1X GlutaMAX, and 100 units/ml of penicillin with 100 µg/ml streptomycin. The episomal iPSC reprogramming plasmids, pCXLE-hOCT3/4, pCXLE-hSK and pCXLE-hMLN were purchased from Addgene. The plasmids used in our experiments were mixed in a ratio of 1:1:1 for efficient reprogramming. 3 μg of expression plasmid mixtures were electroporated into 5×105 HDFs with Amaxa® Nucleofector Kit according to the manufacturer’s instructions. After nucleofection, cells were plated in DMEM containing 10% FCS and 1% penicillin/streptomycin until it reaches 70-80% confluence. The culture medium was replaced the next day by human embryonic stem cell medium (HESM) containing knock-out (KO) DMEM, 20% KO serum replacement (SR), 1 mM NEAAs, 1X GlutaMAX, 0.1 mM β-mercaptoethanol, 1% penicillin/streptomycin and 10ng/ml bFGF (Invitrogen). Between 26-32 days after plating, colonies developed and colonies with a phenotype similar to human ESCs were selected for further cultivation and evaluation. Selected iPSC colonies were mechanically passaged on matrigel (BD, hES qualified matrigel) coated plates containing mTeSR™1 (defined, feeder-free maintenance medium for human ESCs and iPSCs).

### Generation of iPSC-derived neurons

IPSCs were dissociated manually and plated on a non-coated dish in human embryonic stem cell medium (HESM). After 4 days, embryoid bodies (EBs) were formed and transferred to neural differentiation medium containing DMEM/ F12, 1 mM NEAAs, 1× GlutaMAX, 1% penicillin/streptomycin, and 1X N1 supplement (100X) for another 4 days. EBs were plated on matrigel-coated plates for neural rosette formation for 8-10 days with 0.01 mM retinoic acid. Neural rosettes were handpicked and cultured in neural stem cell medium containing DMEM/F12, 1 mM NEAAs, 1X GlutaMAX, 1% penicillin/streptomycin, 1X N1 supplement (100X), 20 ng/mL FGF2 (peprotech), 20 ng/mL EGF (peprotech), and 2 µl/ml B27 supplement (Invitrogen). Terminal neural differentiation was induced by dissociating the neural stem cells (NSCs) using accutase (Sigma) for 20 min at 37°C and plating them on a matrigel-coated plate for attachment. The next day, the medium of these cell cultures were changed to neuronal differentiation medium containing DMEM/F12, 1 mM NEAAs, 1X GlutaMAX, 1% penicillin/streptomycin, 1X N1 supplement (100X), 20 ng/mL BDNF (Peprotech), 20 ng/mL GDNF (Peprotech), 1 mM dibutyryl-cAMP (Sigma) and 2µl/ml B27 supplement (Invitrogen) for 80-90 days.

### Excitatory stimulation of neurons

SCA3 neurons cultured in 6 well plates were washed three times with 2 mL HBSS (balanced salt solution) containing 25 mM Tris, 120 mM NaCl, 15 mM glucose, 5.4 mM KCl, 1.8 mM CaCl_2_, 0.8 mM MgCl_2_, pH 7.4. After treatment with L-glutamate 0.1 mM (Sigma no. G8415) in HBSS for 30 min, cells were washed again three times and left them to recover for 30 min in differentiation media followed by a second 30 min L-glutamate treatment in HBSS, and subsequently cultured in differentiation media or condition media (with or without IGF for 24 and 48 h until analyzed). For analysis of fragmentation and aggregation of Ataxin 3 by western blotting, extracts were analyzed either immediately after lysis or after fractionation.

### Immunocytochemistry

IPSCs-derived neurons from SCA-3 patients were fixated with 4% paraformaldehyde for 20 min. Cells were blocked in 5% normal goat serum and 2% fetal calf serum; subsequently, samples were probed with primary antibodies: TRA-1-81 (Santa Cruz; sc-21706), TRA-2-54 (made by group Peter Andrews lab, The University of Sheffield), OCT-4 (Santa Cruz; sc-5279), SOX-2 (Cell Signaling; #4900S), βIII-tubulin (Abcam; ab7751). Alexa 488 and Cy3-conjugated secondary antibodies were used in combination with Hoechst nuclear staining (1:1000). Images were obtained using a Leica TCS SP8 confocal microscope (Leica Microsystems).

### Statistical analysis

Results were statistically compared using the Student’s *t* test was performed for unpaired or paired groups. A p value of < 0.05 was considered significant (*: *p* < 0.05; **: *p* < 0.01; ***: *p* < 0.001; ****: *p*<0.0001)

## Supplementary Materials

Supplementary Fig. 1. XBP1 deficiency does not alter *Igf1* mRNA levels.

Supplementary Fig. 2 IGF2 expression reduces polyQ79 monomer and aggregation.

Supplementary Fig. 3. IGF2 does not inhibit global protein synthesis or degradation.

Supplementary Fig. 4. Neither proteasome nor autophagy are involved in the reduction of polyQ levels after IGF2 expression.

Supplementary Fig. 5. Controls for secretion experiments.

Supplementary Fig. 6. Gene therapy to deliver IGF2 into the brain.

Supplementary Table 1. Top regulated genes in brain cortex and striatum of XBP1_cKO_ mice.

Supplementary Table 2. List of proteins modified by IGF2 expression.

Supplementary Table 3. Set of proteins modified by IGF2 expression related to actin cytoskeleton dynamics.

Supplementary Table 4. Human post-mortem brain samples.

Supplementary Table 5. Blood samples from HD patients and healthy controls.

Video S1. Neuro2a treated with mock medium.

Video S2. Neuro2a treated with IGF2-enriched medium.

## Acknowledgements.

We are grateful to Dr. Oliver Bracko for providing IGF2 overexpression constructs. We thank the Harvard Brain Tissue Resource Center for providing HD post-mortem samples. We also thank Drs Dimitri Krainc and Katarina Trajkovic for feedback on protocols to detect mHtt secretion. We also thank Dr Cristina Alberini for feedback and providing tools to study IGF2 signaling. We thank Marioly Müller and Dr. Josefina Barrera and her team for their kind help taking the blood samples. We thank Dr. Felipe Oyarzun and Rodrigo Sierpe for the kindly help with the use of the Nanosight NS300. We specially acknowledge Carolina Jerez, Claudia Rivera, Valentina Castillo and Constanza Gonzalez for their technical assistance.

## Authors contribution

P.G-H, P.T-E, C.H and R.L.V designed the study; P.G-H and P.T-E designed, performed and analyzed most of the *in vitro* and *in vivo* experiments; D.W and S.B designed, performed and analyzed the pulse-chase experiments; C.S and F.C analyzed the NTA experiments and assisted with the extracellular vesicle experiments; P.C-C provided the information and blood samples from Chilean HD patients; D.R.H and C.G-B designed and performed the experiments of actin remodeling; P.T, K.A.L and B.J.G performed the microarray analysis using the brains samples from the XBP1_cko_ mice; L.P and L.W performed the quantitative proteomics in Neuro2a cells; C.S and H.G.R performed the Ingenuity pathway analysis; P.S produced the AAV2-IGF2 and AAV2-Mock; G.L.L, N.S and M.S performed the experiments in HEK293 cells using the full-lenghtHttQ_103_-myc plasmid; S.N and L.M.E designed, performed and analyzed the experiments using MSNs derived from HD patients.

## Data and materials availability

MTA may be required for access to AAV-IGF2 and some vectors. All data are available upon request from C.H.

## Supplementary material

**Supplementary Fig. S1.**
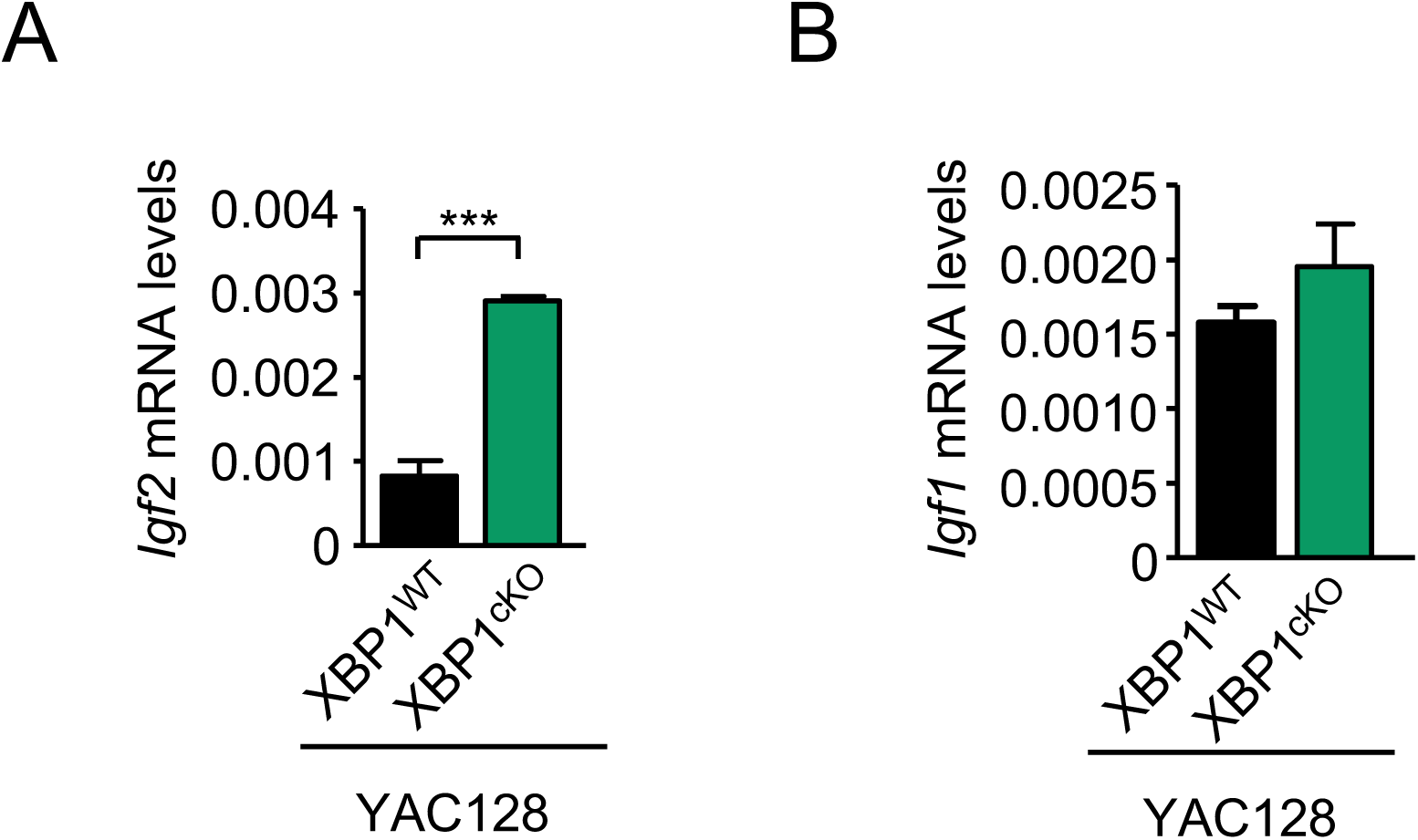
XBP1 deficiency does not alter *Igf1* mRNA levels. **(A)** *Igf2* mRNA levels were measured by real-time PCR in cDNA generated from striatal tissue of 9-months old XBP1_cKO_-YAC128 mice. Data represents the average and SEM of the analysis of 3 animals per group. **(B)** *Igf1* mRNA levels were measured by real-time PCR in cDNA generated from striatal tissue of 9-months old XBP1_cKO_-YAC128 mice. Data represents the average and SEM of 5 animals. Statistically significant differences detected by one-tailed unpaired *t*-test (**: *p* < 0.01).

**Supplementary Fig. S2.**
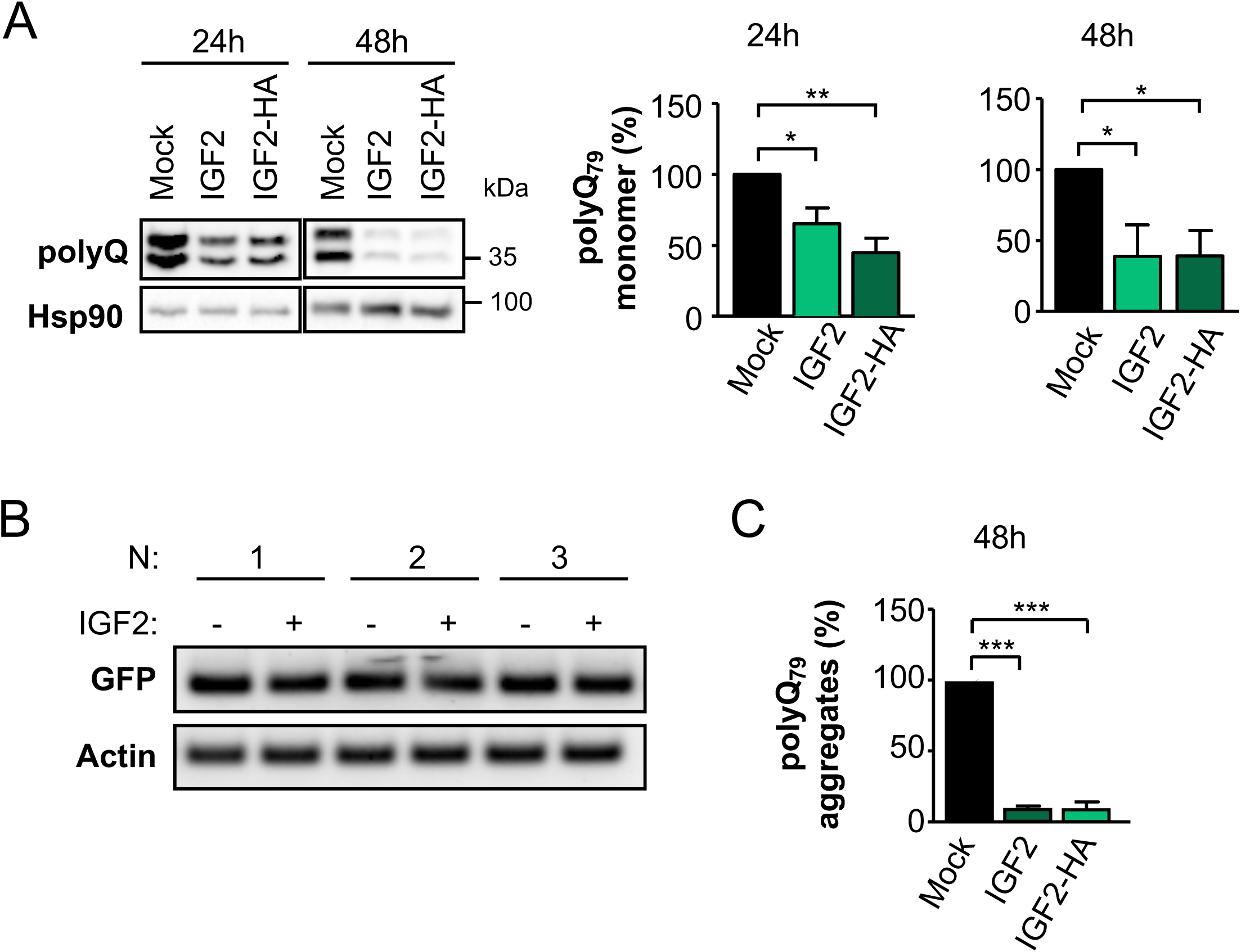
IGF2 expression reduces polyQ_79_ aggregation. **(A)** PolyQ_79_-EGFP monomers from Fig. 2B were analyzed in whole cell extracts by western blot analysis using an anti-GFP antibody (left panel). PolyQ_79_-EGFP monomer levels after 24 and 48h of expression were quantified and normalized to Hsp90 levels (middle and right panel, respectively). **(B)** Neuro2a cells were co-transfected with a polyQ_79_-EGFP expression vector and IGF2 plasmid or empty vector (Mock). 24 h later, cells were collected in Trizol and *Gfp* mRNA levels were measured by semi-quantitative PCR. Values represent the mean and SEM of at least three independent experiments. **c** Quantification of filter trap experiment from Fig. 2c (time point 48 h).

**Supplementary Fig. S3.**
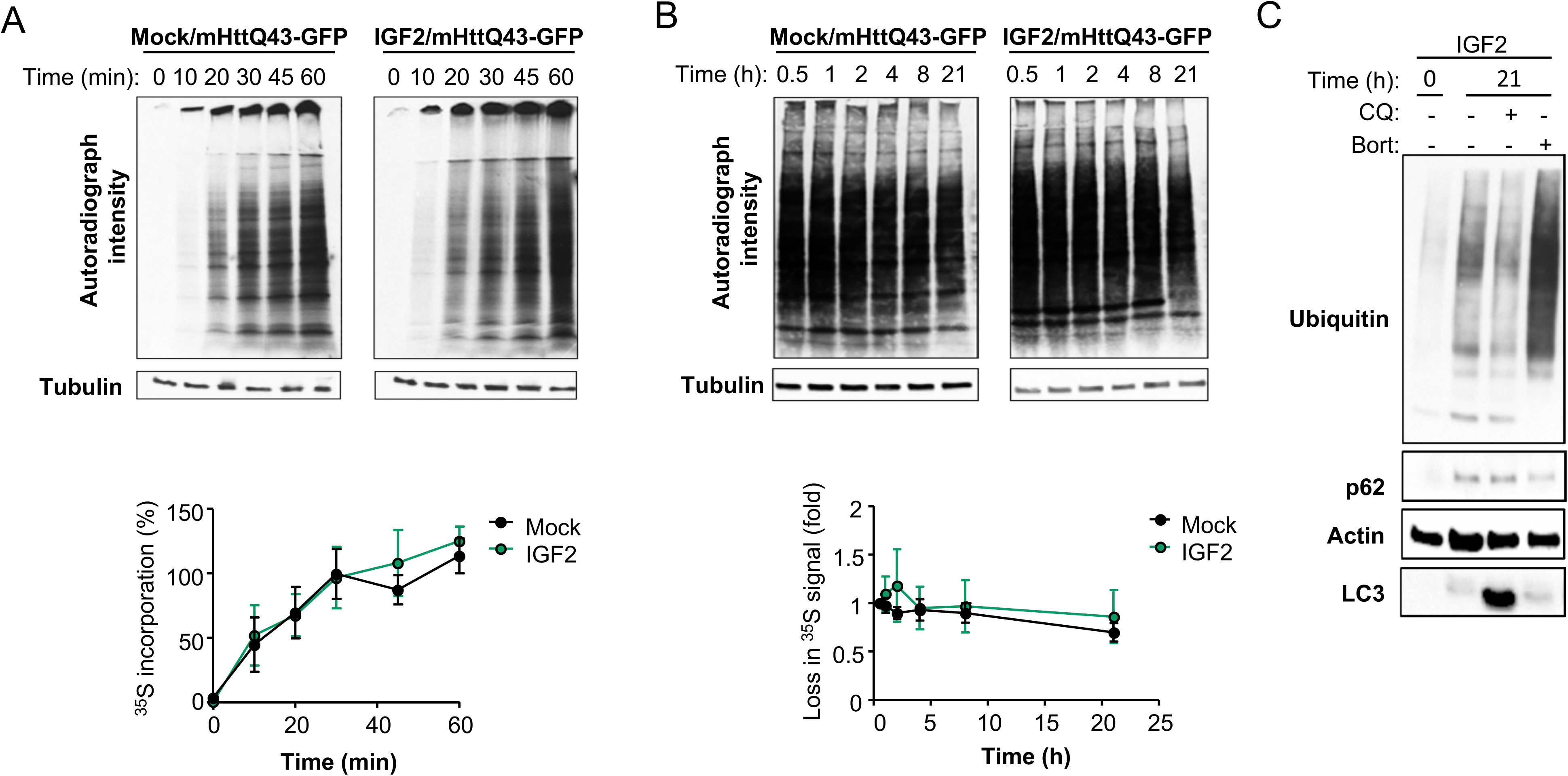
IGF2 does not inhibit global protein synthesis or degradation. **(A)** HEK293T cells were co-transfected with GFP-mHttQ_43_ expression vector with IGF2 plasmid (right, upper panel) or empty vector (left, upper panel), and then pulse labeled with _35_S for indicated time points. Autoradiography (AR) represents total proteins that were synthesized in the time of harvesting cells. All data were normalized to time point 0 h (lower panel). **(B)** HEK293T cells were co-transfected with GFP-mHttQ_43_ expression vector with IGF2 plasmid (right, upper panel) or empty vector (Mock) (left, upper panel), and then pulse labeled with _35_S for indicated time points to follow the decay of the labeled protein while chasing with unlabeled precursor. Autoradiography (AR) represents total extracts in each time (upper panel). All data were normalized to time point 1 h (lower panel). **(C)** HEK293T cells were transfected with GFP-mHttQ_43_ and IGF2 expression vectors. Pulse was performed 24 h after transfection. Cells were treated with 30 μM chloroquine (CQ) or 1 µM bortezomib (Bort) at the beginning of the chasing for additional 21 h (upper panel). Ubiquitin, p62 and LC3 were measured as control of the autophagy inhibition.

**Supplementary Fig. S4.**
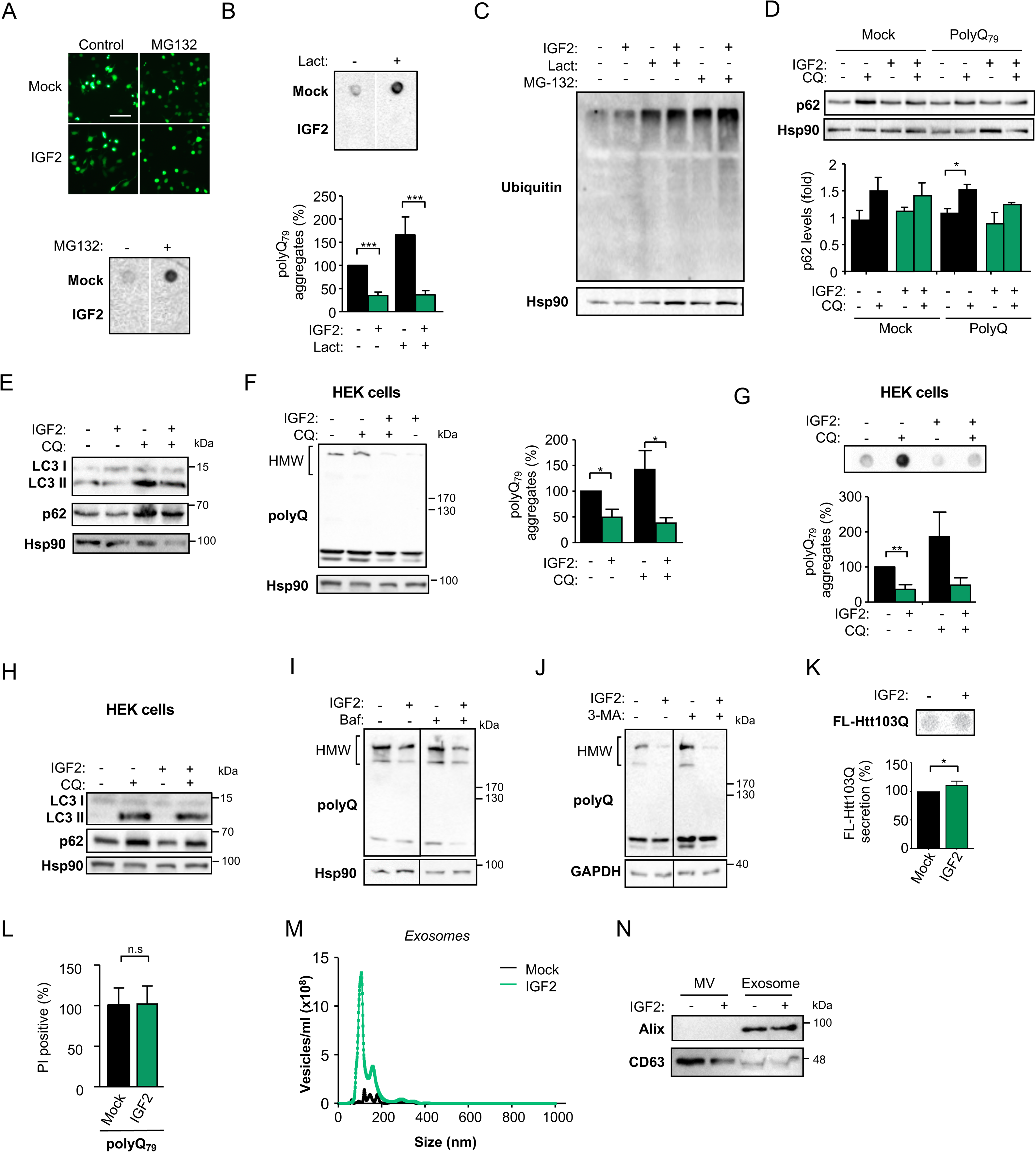
Proteasome, autophagy or conventional secretion are not involved in the reduction of polyQ_79_ levels after IGF2 expression (control experiments). **(A)** Neuro2a cells were co-transfected with expression vectors for polyQ_79_-EGFP and IGF2 or empty vector (Mock). After 8 h, cells were treated with 1 μM MG132 for additional 16 h. PolyQ_79_-EGFP inclusions were visualized by fluorescence microscopy (upper panel) and polyQ_79_-EGFP aggregation levels measured by filter trap (lower panel). Image cropped from the same membrane and film. **(B)** Filter trap assay was performed to detect polyQ_79_-EGFP aggregates using the same cells extracts of Fig. 4E using an anti-GFP antibody (upper panel) and quantified (lower panel). Image cropped from the same membrane and film. **(C)** Neuro2a cells were transfected with IGF2 for 8 h and then treated with 1 μM lactacystin (Lact) or 1 μM MG132 for additional 16 h. Ubiquitin accumulation was measured by western blot to confirm the activity of the inhibitors. Hsp90 expression was measured as loading control. **(D)** Neuro2a cells where co-transfected with IGF2 plasmid or empty vector (Mock) in the presence or absence of polyQ_79_-EGFP expression vector for 8 h, and then treated with 30 μM chloroquine (CQ) for additional 16 h. p62 levels were monitored by western blot using anti-p62 antibody. Hsp90 expression was monitored as loading control (upper panel). p62 levels were quantified and normalized to Hsp90 (lower panel). **(E)** Neuro2a cells were co-transfected with polyQ_79_-EGFP and IGF2 expression vectors or empty vector (Mock) for 8 h, and then treated with 30 μM CQ for additional 16 h. p62 and LC3 levels were measured by western blot as control for autophagy inhibitors in experiment shown in Fig. 4H. Hsp90 levels were determined as loading control. **(F)** HEK293T cells were co-transfected with polyQ_79_-EGFP and IGF2 expression vectors or empty vector (Mock). After 8 h cells were treated with 30 μM chloroquine (CQ) for additional 16 h. PolyQ_79_-EGFP aggregation was analyzed in whole cells extracts by western blot using anti-GFP antibody. Hsp90 expression was monitored as loading control (left panel). PolyQ_79_-EGFP aggregates levels were quantified and normalized to Hsp90 levels (right panel). **(G)** Filter trap assay was performed using the same cells extracts analyzed in (F) and polyQ_79_-EGFP aggregates detected using anti-GFP antibody (upper panel) and quantified (lower panel). **(H)** HEK293T cells were transiently transfected with IGF2 expressing vector (IGF2) or empty plasmid (Mock) for 8 h, and then treated with 30 μM CQ for additional 16 h. p62 and LC3 levels were measured by western blot as control for autophagy inhibitors in experiment shown in Fig. 4F and Fig. 4G. Hsp90 levels were determined as loading control. **(I, J)** Neuro2a cells were co-transfected with polyQ_79_-EGFP and IGF2 or empty vector (Mock) for 8 h and then treated with 200 nM bafilomycin A_1_ (Baf) or 250 mM 3-methyladenine (3-MA) for additional 16 h. PolyQ_79_-EGFP aggregation was analyzed in cell lysates using western blot with an anti-GFP antibody. Hsp90 or GAPDH expressions were monitored as loading control. In all quantifications, average and SEM of at least three independent experiments are shown. Statistically significant differences detected by two-tailed unpaired *t*-test (**: *p* < 0.01; *: *p* < 0.05). **(K)** HEK293 cells were co-transfected with FL-Htt103Q and IGF2 plasmid or empty vector (Mock). 24 h later, culture media was collected and loaded for dot blot detection of FL-Htt103Q using anti-Htt antibody. **(L)** Neuro2a cells were co-transfected with a polyQ_79_-EGFP expression vector and IGF2 plasmid or empty vector (Mock). 48 h later, cell death was measured by FACS after propidium iodide staining. **(M)** NanoSight profile of exosomes purified form IGF2 expressing cells. Isolated exosomes from Neuro2a cells co-transfected with polyQ_79_-EGFP and IGF2 plasmid or empty vector (Mock) were analyzed by NanoSight nanotracking analysis and plotted by size to confirm proper purification method. **(M)** Western blot analysis of enriched fractions of microvesicles (MV) and exosomes using the markers Alix and CD63.

**Supplementary Fig. S5.**
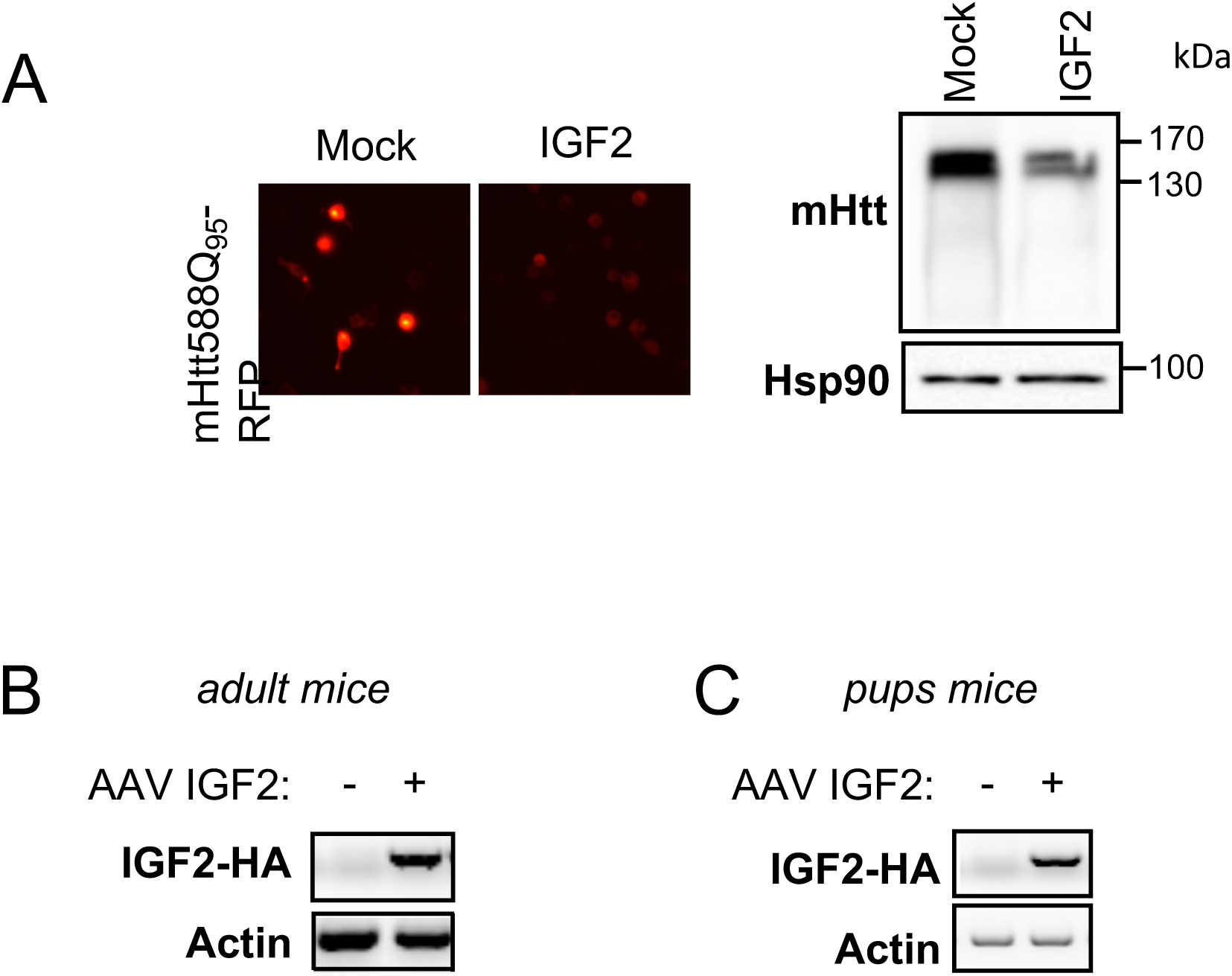
Gene therapy to deliver IGF2 into the brain. (A) Neuro2a cells were co-transfected with expression vectors for mHtt588Q_95_-RFP and IGF2 plasmid or empty vector (Mock) using an AAV back bound. After 24 h, mHttQ_95_-RFP inclusions were visualized by fluorescence microscopy (left panel). Protein aggregation was analyzed in whole cells extracts by western blot (right panel). Hsp90 was measured as loading control. (B) *Igf2-HA* expression was confirmed by conventional PCR in cDNA obtained from dissected striatum of animals co-injected with AAV mHtt588Q_95_-RFP and AAV-IGF2 or AAV-Mock showed in Figure 8A. (C) *Igf2-HA* expression was confirmed by conventional PCR in cDNA obtained from dissected striatum of YAC128 animals injected with AAV-IGF2 or AAV Mock intra-cerebroventricular at neonatal stage P1-P2 showed in Figure 8C.

**Table S1A.** Top regulated genes in brain cortex of XBP1cKO

| Gene | Fold change | q-value (%) |
| --- | --- | --- |
| Klc1 | 1,9 | 10,7 |
| **Acot1** | **1,6** | **0** |
| **Gfap** | **1,6** | **0** |
| **Igf2** | **1,5** | **0** |
| **Mmp14** | **1,5** | **0** |
| **Hopx** | **1,5** | **0** |
| P2ry12 | -1,8 | 84,1 |
| Mpeg1 | -1,6 | 89,2 |
| Pisd-ps3 | -1,6 | 89,2 |
| Kcna6 | -1,6 | 89,2 |
| Slco1c1 | -1,6 | 84,1 |
| Serpina3n | -1,6 | 89,2 |
| Slc25a18 | -1,5 | 89,2 |
| Abhd16a | -1,5 | 106,4 |
| Stx3 | -1,5 | 89,2 |
| Serpina3h | -1,5 | 89,2 |

**Table S1B.** Top regulated genes in brain striatum of XBP1cKO

| Gene | Fold change | q-value (%) |
| --- | --- | --- |
| Prkcd | 1,9 | 103,4 |
| Klc1 | 1,9 | 103,4 |
| Csrp1 | 1,8 | 103,4 |
| **Igf2** | **1,8** | **0** |
| Mff | 1,8 | 103,4 |
| Pdrg1 | 1,7 | 103,4 |
| Igsf1 | 1,6 | 103,4 |
| Fcrls | 1,6 | 103,4 |
| Cpne9 | 1,6 | 103,4 |
| Ramp3 | 1,6 | 103,4 |
| Tbcd | 1,6 | 103,4 |
| Itgb1bp1 | 1,6 | 103,4 |
| Slc13a4 | 1,6 | 103,4 |
| Ppp1r13b | 1,6 | 103,4 |
| **Uhrf1** | **-1,9** | **0** |
| Fam216b | -1,7 | 44,6 |
| P2ry12 | -1,7 | 58,3 |
| Kcna6 | -1,6 | 64,1 |
| Rnd3 | -1,6 | 44,6 |
| Nxph2 | -1,6 | 44,6 |
| Slco1c1 | -1,6 | 64,1 |
| Pbk | -1,6 | 44,6 |
| Prc1 | -1,6 | 44,6 |

**Table S2.**
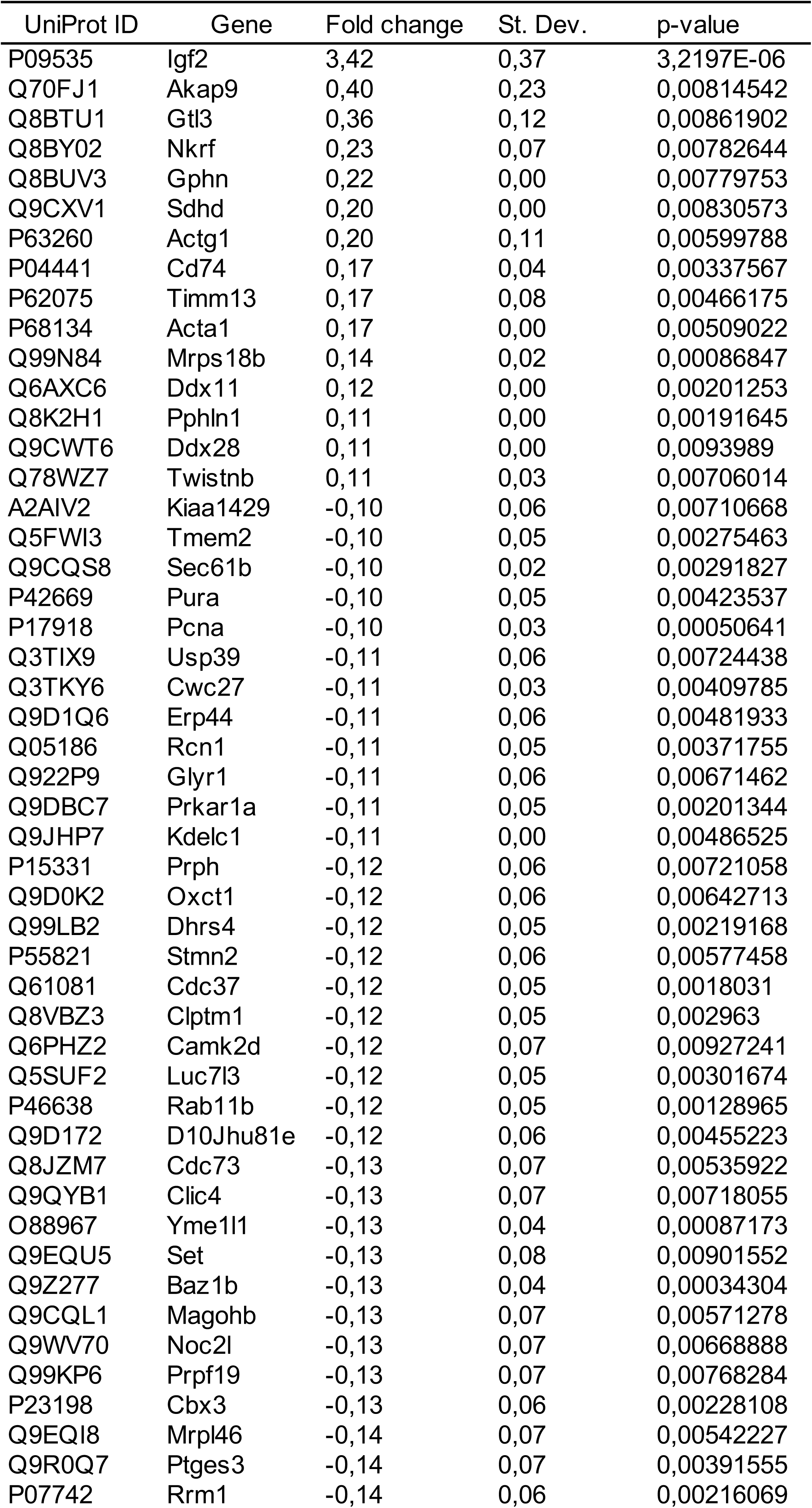
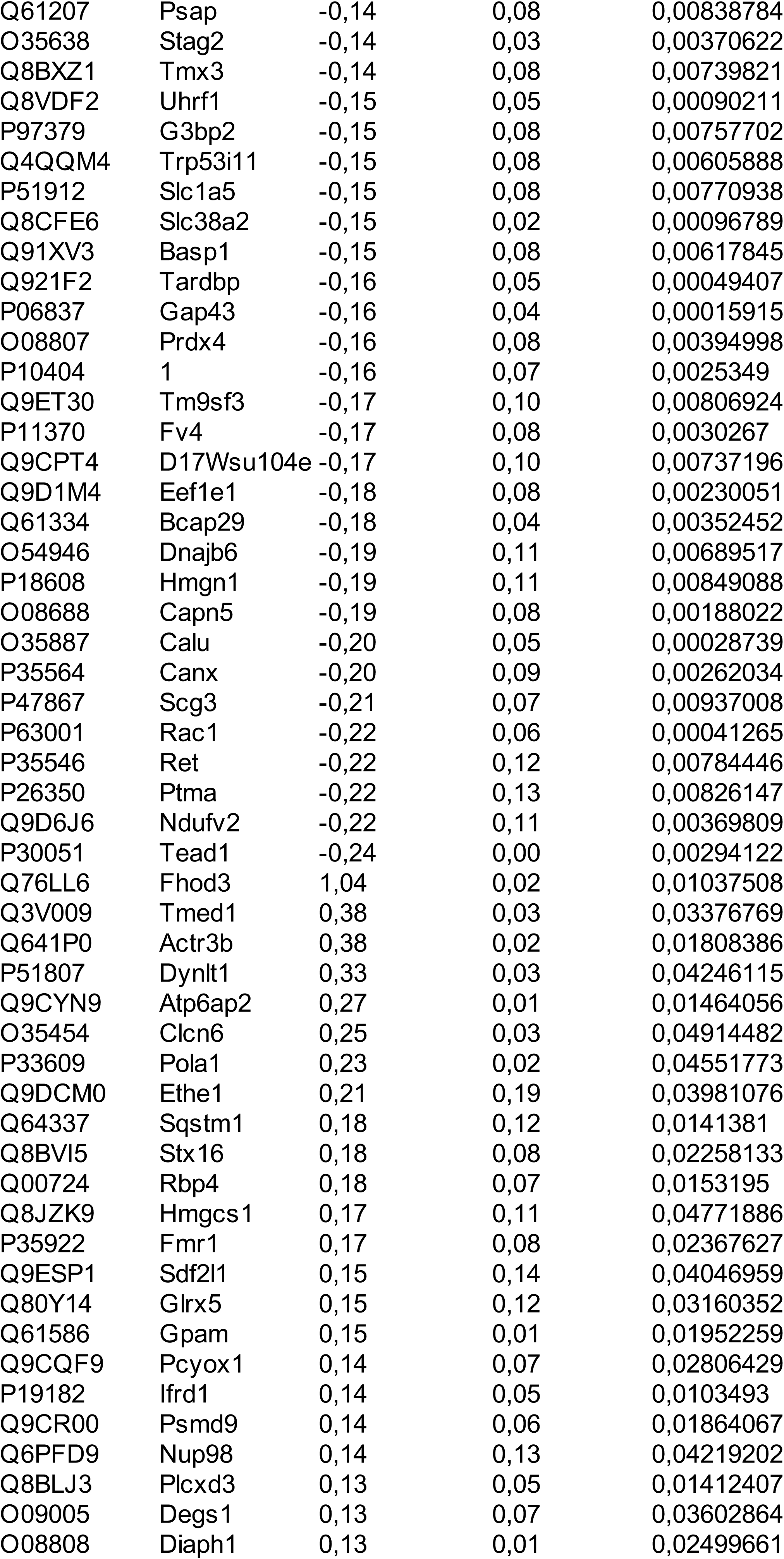
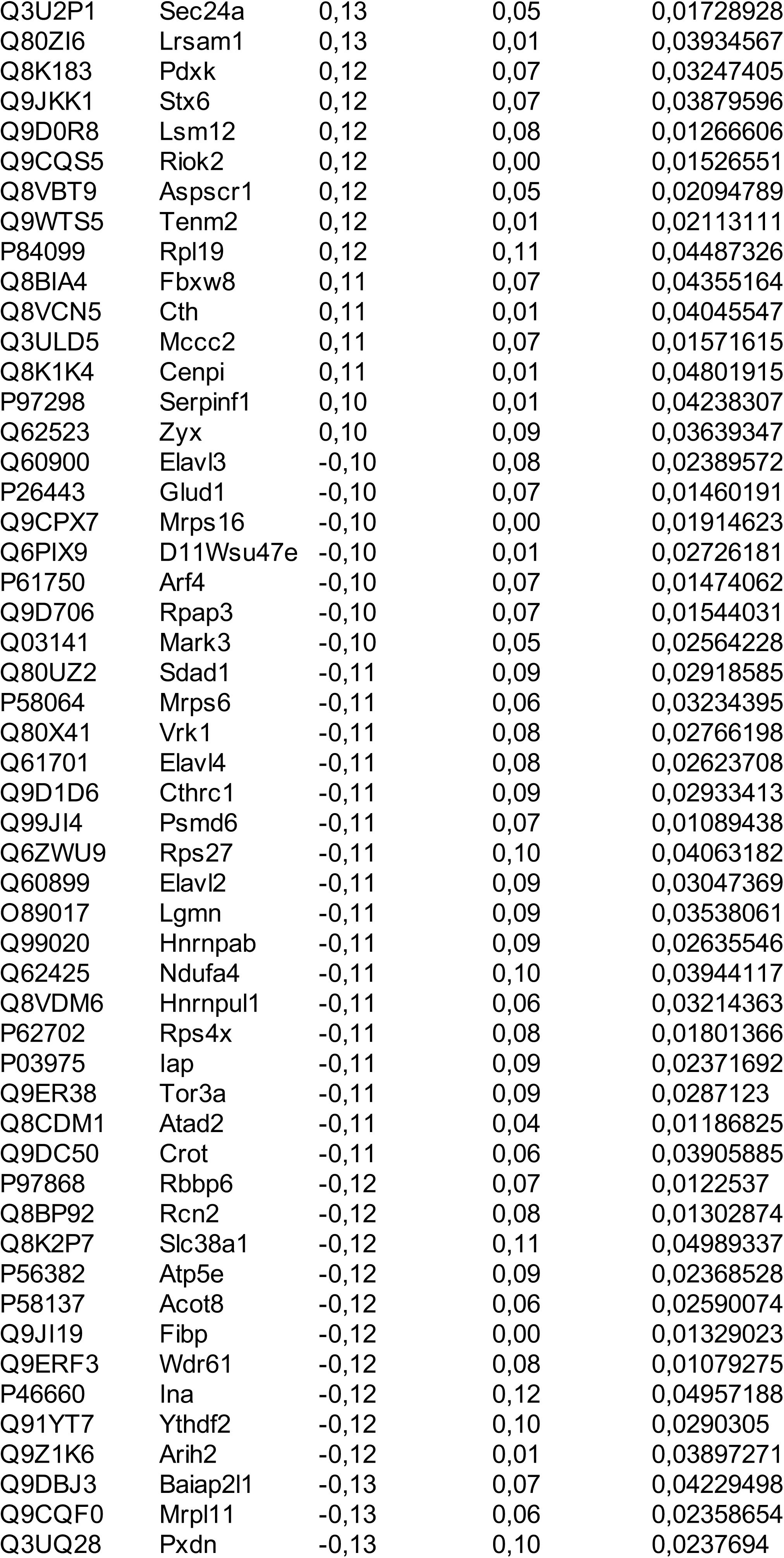
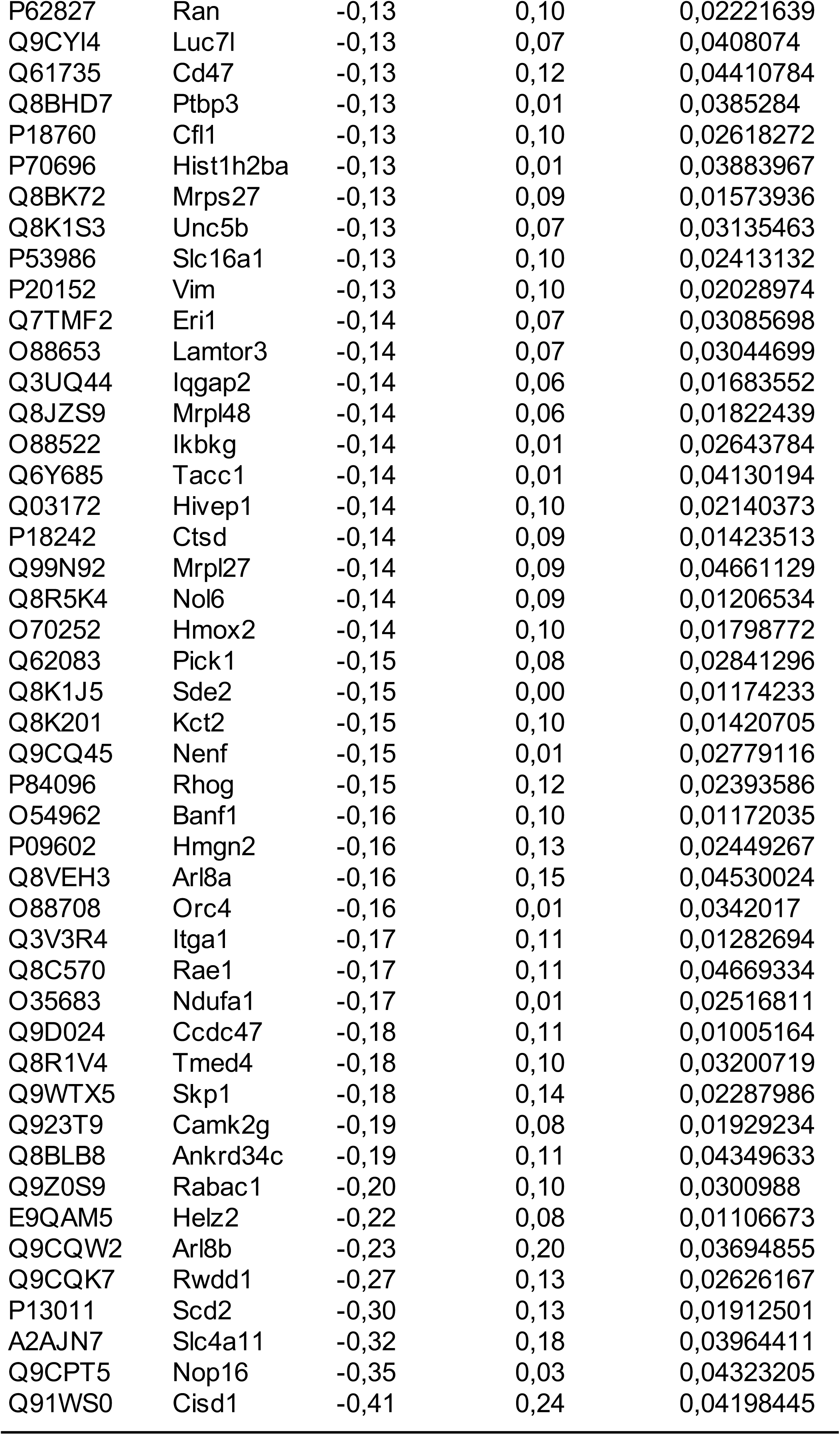
Set of proteins modified by IGF2 expression

**Table S3.** Set of proteins modified by IGF2 expression related to actin cytoskeleton dynamics.

| Gene | Fold change | p-value |
| --- | --- | --- |
| Dbnl | 0,98 | 0,145 |
| Actg1 | 1,15 | 0,006 |
| Acta1 | 1,12 | 0,005 |
| Actr3b | 1,30 | 0,018 |
| Fhod3 | 2,06 | 0,010 |
| Arpc1b | 0,95 | 0,041 |
| Dctn4 | 1,06 | 0,050 |
| Ablim1 | 1,07 | 0,069 |
| Arpc2 | 0,95 | 0,089 |
| Actl6a | 0,96 | 0,123 |
| Rac1 | 0,86 | 0,000 |
| Cdc42 | 0,94 | 0,003 |
| Rhog | 0,90 | 0,024 |
| Arhgap26 | 1,02 | 0,040 |
| Cfl1 | 0,91 | 0,026 |
| Iqgap2 | 0,91 | 0,017 |
| Zyx | 1,07 | 0,036 |
| BAIAP2L1 | 0,92 | 0,042 |
| DIAPH1 | 1,09 | 0,025 |
| TACC1 | 0,91 | 0,041 |
| PICK1 | 0,90 | 0,028 |
| Vim | 0,91 | 0,020 |
| Ran | 0,92 | 0,022 |
| Akap9 | 1,32 | 0,008 |
| Dnajb6 | 0,88 | 0,007 |
| STMN2 | 0,92 | 0,006 |
| CLIC4 | 0,92 | 0,007 |
| Dynlt1 | 1,26 | 0,042 |
| Sdad1 | 0,93 | 0,029 |
| Ina | 0,92 | 0,050 |
| Slc16a1 | 0,91 | 0,024 |
| Prkar1a | 0,93 | 0,002 |
| PRPH | 0,92 | 0,007 |

**Table S4.** Human post-mortem brain samples.

| Group | Sex | Age | PMI | Region | Familiar antecedents |
| --- | --- | --- | --- | --- | --- |
| Control | m | 53 | 15,98 | caudate | 0 |
| Control | m | 63 | 17,92 | caudate | 0 |
| Control | m | 64 | 18,85 | putamen | 0 |
| Control | m | 55 | 19,18 | caudate | 0 |
| Control | f | 58 | 19,38 | putamen | 0 |
| Control | m | 54 | 19,9 | putamen | 0 |
| Control | m | 63 | 19,91 | caudate | 0 |
| Control | m | 63 | 20,58 | caudate | 0 |
| Control | m | 65 | 21 | caudate | 0 |
| Control | f | 60 | 28,15 | caudate | 0 |
| Control | m | 53 | 23,57 | caudate | 0 |
| Control | m | 58 | 26,3 | caudate | 0 |
| Control | m | 52 | 19,55 | caudate | 0 |
| HD, Grade 3 | f | 51 | 14,8 | caudate | 1 |
| HD, Grade 3 | m | 60 | 14,9 | putamen | N/a |
| HD, Grade 3 | f | 58 | 15,33 | putamen | 1 |
| HD, Grade 3 | m | 39 | 16,08 | putamen | 1 |
| HD, Grade 3 | m | 52 | 16,33 | caudate | 1 |
| HD, Grade 3 | m | 65 | 19,9 | caudate | N/a |
| HD, Grade 3 | m | 38 | 25,92 | caudate | N/a |
| HD, Grade 3 | m | 62 | 19,4 | caudate/putamen | N/a |
| HD, Grade 3 | f | 64 | 19,65 | caudate | 1 |
| HD, Grade 4 | f | 59 | 12,91 | putamen | 1 |
| HD, Grade 4 | m | 48 | 19,68 | caudate | 1 |
| HD, Grade 4 | m | 48 | 27,15 | caudate | 1 |
| HD, Grade 4 | m | 58 | 27,32 | putamen | 1 |

**Table S5.** Blood samples from HD chilean patients and healthy controls.

| Age | Familiar history | Participant category | Allele 1 | Allele 2 | Onset year | UHDRS Total Funtional Capacity |
| --- | --- | --- | --- | --- | --- | --- |
| 53 | 1 | pre-manifest HD |  |  |  | 13 |
| 67 | 1 | pre-manifest HD |  |  |  | 13 |
| 23 | 1 | pre-manifest HD | 18 | 50 |  | 13 |
| 77 | 1 | pre-manifest HD |  |  |  | 13 |
| 25 | 1 | pre-manifest HD |  |  |  | 13 |
| 48 | 1 | pre-manifest HD |  |  |  | 13 |
| 35 | 1 | pre-manifest HD |  |  |  | 13 |
| - | - | pre-manifest HD |  |  |  | 13 |
| 40 | 1 | HD | 26 | 47 | 2011 | 12 |
| 53 | 1 | HD | 21 | 43 | 2014 | 7 |
| 33 | 1 | HD |  |  | 2012 | 8 |
| 63 | 1 | HD | 17 | 40 | 2011 | 13 |
| 60 | 1 | HD |  |  | 2015 | 6 |
| 39 | 0 | HD |  |  | 2011 | 11 |
| 37 | 1 | HD | 15 | 48 | 2014 | 10 |
| 61 | 1 | HD |  |  | 2015 | 10 |
| 55 | 1 | HD |  |  | 2008 | 2 |
| 43 | 1 | HD |  |  | 2008 | 6 |
| 45 | 0 | HD |  |  | 2006 | 4 |
| 63 | 1 | HD |  |  | 2010 | 10 |
| 49 | 1 | HD | 17 | 42 | 2006 | 10 |
| 50 | 1 | HD |  |  | 2006 | 6 |

